# A heterochromatin-specific RNA export pathway facilitates piRNA production

**DOI:** 10.1101/596171

**Authors:** Mostafa F. ElMaghraby, Peter Refsing Andersen, Florian Pühringer, Katharina Meixner, Thomas Lendl, Laszlo Tirian, Julius Brennecke

## Abstract

PIWI-interacting RNAs (piRNAs) guide transposon silencing in animals. The 22-30nt piRNAs are processed in the cytoplasm from long non-coding RNAs. How piRNA precursors, which often lack RNA processing hallmarks of export-competent transcripts, achieve nuclear export is unknown. Here, we uncover the RNA export pathway specific for piRNA precursors in the *Drosophila* germline. This pathway requires Nxf3-Nxt1, a variant of the hetero-dimeric mRNA export receptor Nxf1-Nxt1. Nxf3 interacts with UAP56, a nuclear RNA helicase essential for mRNA export, and CG13741/Bootlegger, which recruits Nxf3-Nxt1 and UAP56 to heterochromatic piRNA source loci. Upon RNA cargo binding, Nxf3 achieves nuclear export via the exportin Crm1, and accumulates together with Bootlegger in peri-nuclear nuage, suggesting that after export, Nxf3-Bootlegger delivers precursor transcripts to the piRNA processing sites. Our findings indicate that the piRNA pathway bypasses nuclear RNA surveillance systems to achieve export of heterochromatic, unprocessed transcripts to the cytoplasm, a strategy also exploited by retroviruses.

## INTRODUCTION

Integrity of the germline genome is essential for the survival of multicellular species. In animal gonads, a class of small regulatory RNAs called PIWI-interacting RNAs (piRNAs) guide Argonaute effector proteins to silence transposable elements, thereby counteracting their mutagenic impact (Czech et al., 2018; Ozata et al., 2018; Siomi et al., 2011). piRNAs are processed from single-stranded precursors, which are transcribed by RNA polymerase II from transposon-rich loci termed piRNA clusters. As piRNA biogenesis occurs in peri-nuclear processing centers, the long non-coding precursor transcripts must be exported from the nucleus to the cytoplasm.

mRNAs, the most prominent RNA polymerase II transcripts, are exported through nuclear pore complexes primarily by the Nuclear RNA export factor 1 (Nxf1/Tap) and its binding partner Nxt1/p15 (Cullen, 2000; Kohler and Hurt, 2007; Reed and Hurt, 2002). The Nxf1-Nxt1 heterodimer is recruited to export-competent mRNAs after successful completion of RNA-processing events such as 5′ capping, splicing, and 3′ end-formation (Heath et al., 2016). Nuclear mRNA surveillance systems ensure that unprocessed transcripts are not exported, but instead are degraded within the nucleus. mRNA processing is therefore a critical step in licensing export of RNA polymerase II transcripts to the cytoplasm.

In *Drosophila*, only few piRNA precursors are spliced and poly-adenylated, and these require the mRNA export receptor Nxf1-Nxt1 for nuclear exit (Dennis et al., 2016; Goriaux et al., 2014; Mohn et al., 2014). Most piRNA precursors instead lack RNA processing marks characteristic for mRNAs. These precursors originate from heterochromatic loci, whose expression depends on Rhino, a variant of the conserved heterochromatin protein 1 (HP1) (Klattenhoff et al., 2009). Rhino, via its adaptor protein Deadlock, recruits effector proteins that stimulate transcription initiation within heterochromatin (through the TFIIA-L homolog Moonshiner) (Andersen et al., 2017), and that suppress co-transcriptional RNA processing events such as splicing or 3′ cleavage and polyadenylation (through the Rai1 homolog Cutoff) (Chen et al., 2016; Mohn et al., 2014; Zhang et al., 2014). Rather than imposing transcriptional silencing similar to HP1, Rhino thereby facilitates non-canonical transcription of heterochromatic piRNA source loci to allow the production of non-coding precursors for transposon-targeting piRNAs. How the resulting unprocessed piRNA precursors escape nuclear RNA quality control, and how they are transported to cytoplasmic piRNA processing centers are central open questions.

Here, we identify a nuclear export pathway dedicated to piRNA precursors. It involves Nuclear export factor 3 (Nxf3), which is required for piRNA production, transposon silencing and fertility, and which evolved from the principal mRNA-export receptor Nxf1. Instead of relying on RNA processing events for recruitment, Nxf3 is targeted to nascent piRNA precursors via Rhino, Deadlock, and CG13741/Bootlegger, a previously unknown effector protein of Rhino-dependent heterochromatin expression. After cargo-binding, Nxf3 mediates piRNA precursor export in a Crm1/Xpo1-dependent manner. In the cytoplasm, Nxf3, Bootlegger and piRNA precursors accumulate in peri-nuclear nuage where piRNA biogenesis occurs. This dedicated RNA export pathway—formed by Nxf1 duplication and neo-functionalization—therefore bypasses nuclear RNA surveillance systems and transports piRNA precursors from their heterochromatic origins to their cytoplasmic processing sites.

## RESULTS

### Nxf3 localizes to piRNA transcription and processing sites

To visualize the transport route of Rhino-dependent piRNA precursors in ovaries, we performed fluorescent *in situ* hybridization (RNA FISH) experiments directed against transcripts from piRNA *cluster42AB*, the most prominent piRNA source locus genome-wide (Brennecke et al., 2007). *cluster42AB* transcripts were enriched in nuclear foci as well as in cytoplasmic foci lining the nuclear envelope (Figure 1a, Figure S1a). The nuclear foci correspond to piRNA clusters where Rhino licenses piRNA precursor transcription (Figure 1b) (Klattenhoff et al., 2009; Mohn et al., 2014; Zhang et al., 2014). The peri-nuclear foci in the cytoplasm mark ‘nuage’, electron-dense RNP granules where Vasa and other factors orchestrate piRNA biogenesis (Figure 1c) (Lim and Kai, 2007; Mahowald, 1971; Malone et al., 2009; Zhang et al., 2012).

**Figure 1.**
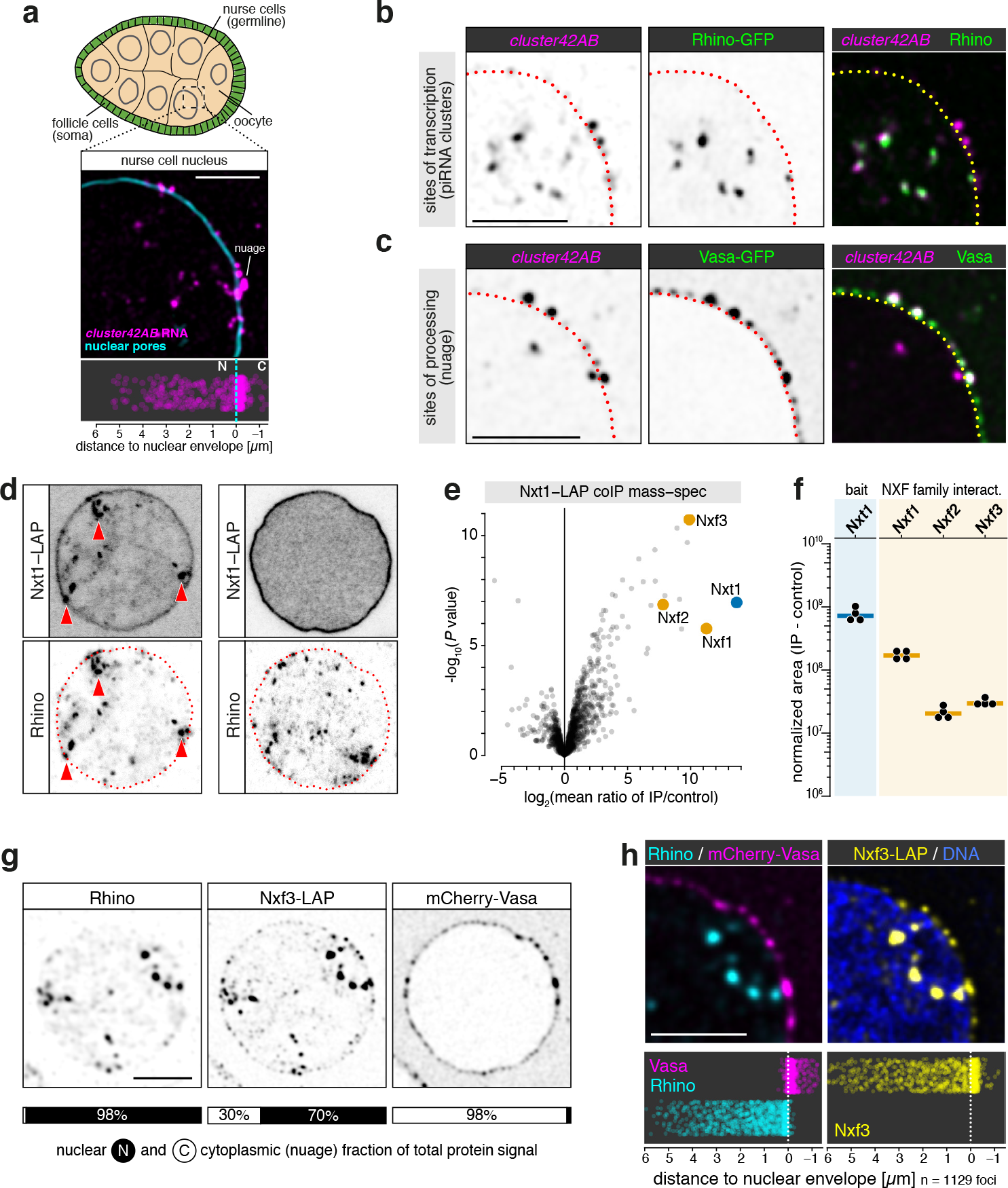
Nxf3 localizes to piRNA clusters and nuage. **a**, Top: cartoon of *Drosophila* stage 7 egg chamber; nurse cells (germline) in beige, somatic support cells in green. Middle: confocal image (scale bar: 5 µm) showing part of a nurse cell nucleus; nuclear pores stained with wheat germ agglutinin (cyan); piRNA *cluster42AB* precursor in magenta (RNA FISH;). Bottom: jitter plot showing positions of *cluster42AB* RNA FISH foci relative to nuclear envelope (n = 509 from 11 nuclei). **b, c**, Confocal images (scale bars: 5 µm) showing *cluster42AB* transcripts and GFP-Rhino (**b**) or GFP-Vasa (**c**) in nurse cells (yellow dotted line: center of nuclear envelope; see Figure S1a). **d**, Localization of Nxt1-LAP or Nxf1-LAP (top panels) alongside Rhino (bottom panels) in nurse cells (red dotted line: nuclear envelope). **e**, Volcano plot showing enrichment values and corresponding significance levels for proteins co-purifying with Nxt1-LAP versus control (nuclear LAP) from ovary lysates (n = 3). **f**, Peptide peak intensities (experiment minus respective replicate control) for the Nxt1-LAP bait and the protein interactors marked in **f** (bars: median values). **g**, Inverted black/white confocal image of a nurse cell nucleus (scale bar: 5 µm) showing localization of Rhino, mCherry-Vasa, and Nxf3-LAP. Cartoons below indicate sum of nuclear/cytoplasmic signal intensities of >1000 foci per protein. **h**, Part of nucleus (scale bar: 5 µm) shown in (**g)** indicating localization of Nxf3-LAP (yellow), Rhino (cyan), and mCherry-Vasa (magenta). Jitter plots below show positions of indicated protein foci relative to the nuclear envelope (>1000 foci per protein).

Export of mRNA from the nucleus requires the association of the heterodimeric nuclear RNA export receptor Nxf1-Nxt1 (Tap-p15) with mRNA cargo, which is enabled by the nuclear RNA helicase UAP56/Hel25E and the associated THO complex, present from yeast to human (Heath et al., 2016; Kohler and Hurt, 2007; Luo et al., 2001; Strasser and Hurt, 2001). In the *Drosophila* germline, both, UAP56 and the THO complex are enriched at Rhino-dependent piRNA clusters and are required for efficient piRNA production from these loci (Hur et al., 2016; Zhang et al., 2012; Zhang et al., 2018). To ask whether Nxf1-Nxt1 might also export germline piRNA precursors, we generated flies expressing Nxf1 or Nxt1 with a ‘localization and affinity-purification’ (LAP) tag consisting of GFP and a triple FLAG peptide. Nxt1-LAP was enriched at nuclear Rhino foci (Figure 1d). Nxf1-LAP instead was uniformly distributed in the nucleus, with enrichment at the nuclear envelope, as also seen for Nxt1-LAP (Figure 1d). This suggested that the mRNA export co-factor Nxt1 functions at Rhino-dependent piRNA clusters, probably in an Nxf1-independent manner.

To identify proteins acting together with Nxt1 at piRNA clusters, we affinity-purified Nxt1-LAP from ovary lysates and analyzed co-eluting proteins by quantitative mass spectrometry. In addition to Nxf1, the NXF variants Nxf2 and Nxf3 were highly enriched in Nxt1-LAP eluates (Figure 1e), albeit with lower peptide levels (Figure 1f). We confirmed these findings using an independent fly line, expressing HA-tdTomato-tagged Nxt1 from the endogenous locus (Figure S1b, c). Nxf2 and Nxf3 are expressed predominantly in ovaries (Figure S1d) (Brown et al., 2014), and were genetically identified as putative piRNA pathway factors (Czech et al., 2013). As Nxf2 acts in piRNA-guided heterochromatin formation, and not in nuclear mRNA or piRNA precursor export (J. Batki, J. Schnabl, J. Brennecke; manuscript in preparation), we focused on Nxf3, an orphan NXF variant with unknown function (Herold et al., 2001; Herold et al., 2000; Herold et al., 2003).

Tagging the endogenous *nxf3* locus with a C-terminal LAP-tag revealed Nxf3 expression specifically in nurse cells (Figure S1e). Its nuclear localization strongly overlapped with that of Rhino (Figure 1g, h), indicating that Nxf3 functions at piRNA clusters together with Nxt1. However, unlike the strictly nuclear Rhino protein, Nxf3-LAP was also enriched in cytoplasmic foci lining the nuclear envelope (Figure 1g, h). All of these cytoplasmic Nxf3 foci were positive for the piRNA biogenesis factor Vasa, indicating that they represent nuage granules. Indeed, immuno-gold electron microscopy revealed localization of Nxf3-LAP within peri-nuclear, electron-dense nuage compartments, as well as in adjacent nuclear regions (Figure S1f, g) where Rhino is often observed (Zhang et al., 2012).

Of all known proteins localizing to Rhino-dependent piRNA clusters, Nxf3 is unique in that it is also enriched in nuage. Considering this dual localization, together with Nxf3’s homology to the mRNA export receptor Nxf1 (Herold et al., 2000), we hypothesized that Nxf3 is recruited to Rhino-dependent piRNA clusters from where it transports piRNA precursors through nuclear pore complexes to cytoplasmic piRNA processing centers.

### piRNA precursor export requires Nxf3

Our model predicts that Nxf3 is required for efficient mature piRNA production. To test this, we generated flies carrying *nxf3* null alleles (Figure S2a, b). Based on small RNA sequencing, ovaries lacking Nxf3 exhibited ~5–10-fold reduced levels of piRNAs originating from Rhino-dependent piRNA source loci (Figure 2a-c). In contrast, piRNAs originating from the Rhino-independent piRNA source loci *cluster20A* or *flamenco* (Klattenhoff et al., 2009; Mohn et al., 2014) were not reduced in *nxf3* mutant ovaries (Figure 2b, c). In line with this, Rhino-independent piRNA precursors are spliced, poly-adenylated, and exported by Nxf1-Nxt1 (Dennis et al., 2016; Goriaux et al., 2014; Mohn et al., 2014).

**Figure 2.**
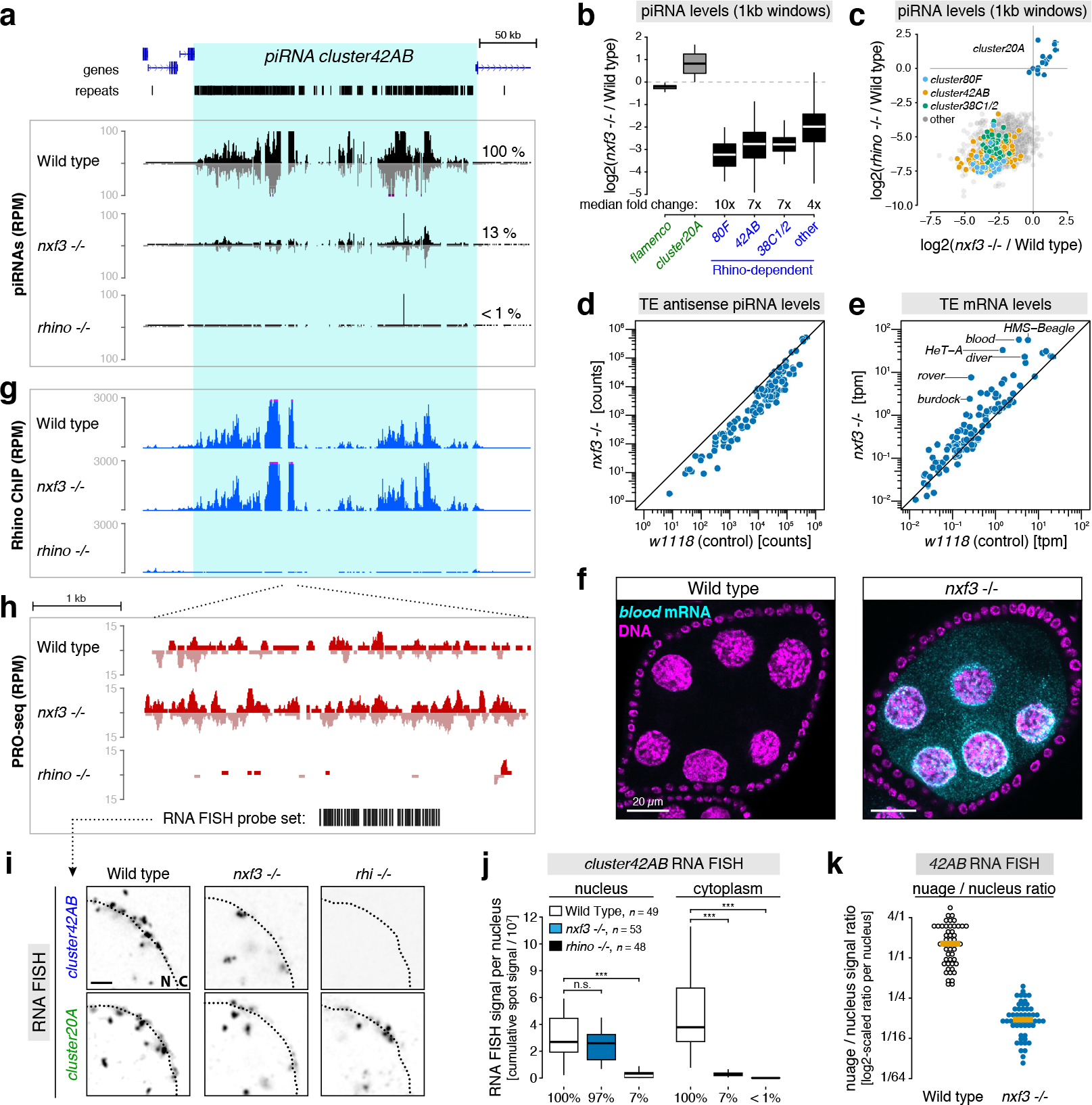
Nxf3 is required for piRNA precursor export. **a**, UCSC genome browser panel showing ovarian piRNA levels (reads per million miRNA reads) at *cluster42AB* in indicated genotypes. **b**, Boxplot showing changes in ovarian piRNA levels for indicated piRNA clusters (split into 1kb tiles) in *nxf3* mutants, relative to control. Box plot displays median (line), first and third quartiles (box), and highest/lowest value within 1.5 × interquartile range (whiskers). **c**, Scatter plot displaying log2-fold changes of piRNAs mapping uniquely to 1kb tiles from Rhino-dependent piRNA clusters in *rhino* versus *nxf3* mutants (relative to control). Tiles from major piRNA clusters are colored, *cluster20A* tiles serve as Rhino-independent control group. **d, e**, Scatter plots displaying levels of antisense piRNAs (**d**) or levels of sense mRNA reads (**e**) mapping to consensus transposon sequences from control versus *nxf3* mutant ovary samples. **f**, Confocal images of wildtype (left) and *nxf3* mutant (right) egg chambers showing *blood* transposon mRNA (FISH: cyan; DAPI stain: magenta). **g**, UCSC genome browser panel showing Rhino occupancy (ChIP-seq; reads per million reads) at *cluster42AB* in indicated genotypes. **h**, 5kb part of *cluster42AB* with PRO-seq signal from the indicated genotypes in red. Black bars indicate the 48 oligo FISH probes used in panel (**i**). **i**, Representative confocal images of parts of germline nuclei (genotypes indicated) stained with RNA-FISH for *cluster42AB* transcripts (Rhino dependent) and *cluster20A* transcripts (Rhino-independent). **j**, Quantification of *cluster42AB* RNA FISH signal in nurse cells with indicated genotype (boxplot definition as in **b**; ***P < 0.0001 based on Mann–Whitney–Wilcoxon test; number of analyzed germline nuclei indicated). **k**, Ratio of total *cluster42AB* RNA FISH signal in nuage versus nucleus quantified per nurse cell nucleus (genotypes indicated; orange bars: median values).

For many germline-active transposons, the loss of piRNAs from Rhino-dependent source loci resulted in ~10-fold reduced levels of antisense, silencing-competent piRNAs (Figure 2d, Figure S2c, d). Consistent with this, several transposons were de-repressed in *nxf3* mutant ovaries (Figure 2e, Figure S2e). Among them are *blood* (Figure 2f, Figure S2f), *HMS*-*Beagle, rover, diver*, and *HeT-A*, retro-transposons whose de-silencing has been linked to severe oocyte DNA damage in piRNA pathway mutants (Durdevic et al., 2018; Senti et al., 2015; Wang et al., 2018). Indeed, *nxf3* mutants were female sterile, despite globally unperturbed levels of ovarian mRNAs (Figure S2g, h).

To determine how Nxf3 loss impairs piRNA production, we analyzed whether Rhino-dependent piRNA source loci are still specified and transcribed in *nxf3* mutants. Rhino’s genome-wide enrichment at piRNA clusters was unchanged in *nxf3* mutant ovaries (Figure 2g; Figure S2i), as was the localization of factors co-localizing with Rhino at piRNA clusters (e.g. Deadlock, Cutoff, Moonshiner, UAP56; Figure S3a, b) (Andersen et al., 2017; Mohn et al., 2014; Pane et al., 2011; Wehr et al., 2006; Zhang et al., 2012; Zhang et al., 2014). piRNA cluster transcription, as determined by global precision nuclear run-on sequencing (PRO-seq) (Kwak et al., 2013), was also virtually unchanged in *nxf3* mutant ovaries (Figure 2h; Figure S2i). The observed piRNA losses in *nxf3* mutants must therefore be due to defects downstream of piRNA cluster specification and transcription. To assay the fate of piRNA precursors in *nxf3* mutants, we determined the subcellular localization of *cluster42AB* transcripts, with the Rhino-independent *cluster20A* serving as a control. In *rhino* mutants, nuclear and cytoplasmic *cluster42AB* foci (marking transcription and piRNA biogenesis sites, respectively) were lost (Figure 2i, j). The nuclear *cluster42AB* foci were unaffected in *nxf3* mutants (Figure 2i, j), consistent with the PRO-seq data (Figure 2h), whereas loss of Nxf3 substantially reduced the cytoplasmic *cluster42AB* signal (Figure 2i-k). By contrast, nuclear and cytoplasmic accumulation of Rhino-independent *cluster20A* transcripts was unaffected in *nxf3* mutants (Figure 2i; Figure S3c, d). As *cluster42AB* transcripts did not accumulate in *nxf3* mutant nurse cell nuclei, our findings indicate that piRNA precursors, which cannot be exported in the absence of Nxf3 are instead targeted for degradation in the nucleus. Consistent with this, we observed a ~2-3-fold reduction in steady-state RNA levels from Rhino-dependent piRNA clusters (Figure S3e, f).Altogether, our data support a direct role for Nxf3 in stabilization and nuclear export of Rhino-dependent piRNA cluster transcripts.

### Nxf3 binds piRNA precursors

Nxf3-LAP extensively co-localized with *cluster42AB* transcripts in the nucleus and nuage (Figure 3a). Supporting a direct interaction between Nxf3 and RNA, electrophoretic mobility shift assays revealed that Nxf3’s N-terminal RRM-LRR domain (RNA recognition motif and Leucine-rich repeat) is capable of RNA binding *in vitro* (Figure S4a, b). To test whether Nxf3 associates with Rhino-dependent piRNA precursors *in vivo*, we performed RNA immunoprecipitation (RIP) experiments on Nxf3-LAP expressing ovaries, ovaries expressing UAP56-LAP (which binds Rhino-dependent piRNA precursors) (Zhang et al., 2012; Zhang et al., 2018), and ovaries expressing nuclear LAP alone. Based on quantitative RT-PCR, Nxf3-LAP RIP eluates recovered ~10% of *cluster42AB* transcripts present in the lysate (input) (Figure 3b). This recovery was in a similar range compared to UAP56-LAP RIP and several hundred-fold more than in the LAP control eluate. In contrast, transcripts from the Rhino-independent piRNA *cluster20A*, abundant house-keeping mRNAs, or rRNAs were not enriched above background in Nxf3-LAP RIP eluates (Figure 3b).

**Figure 3.**
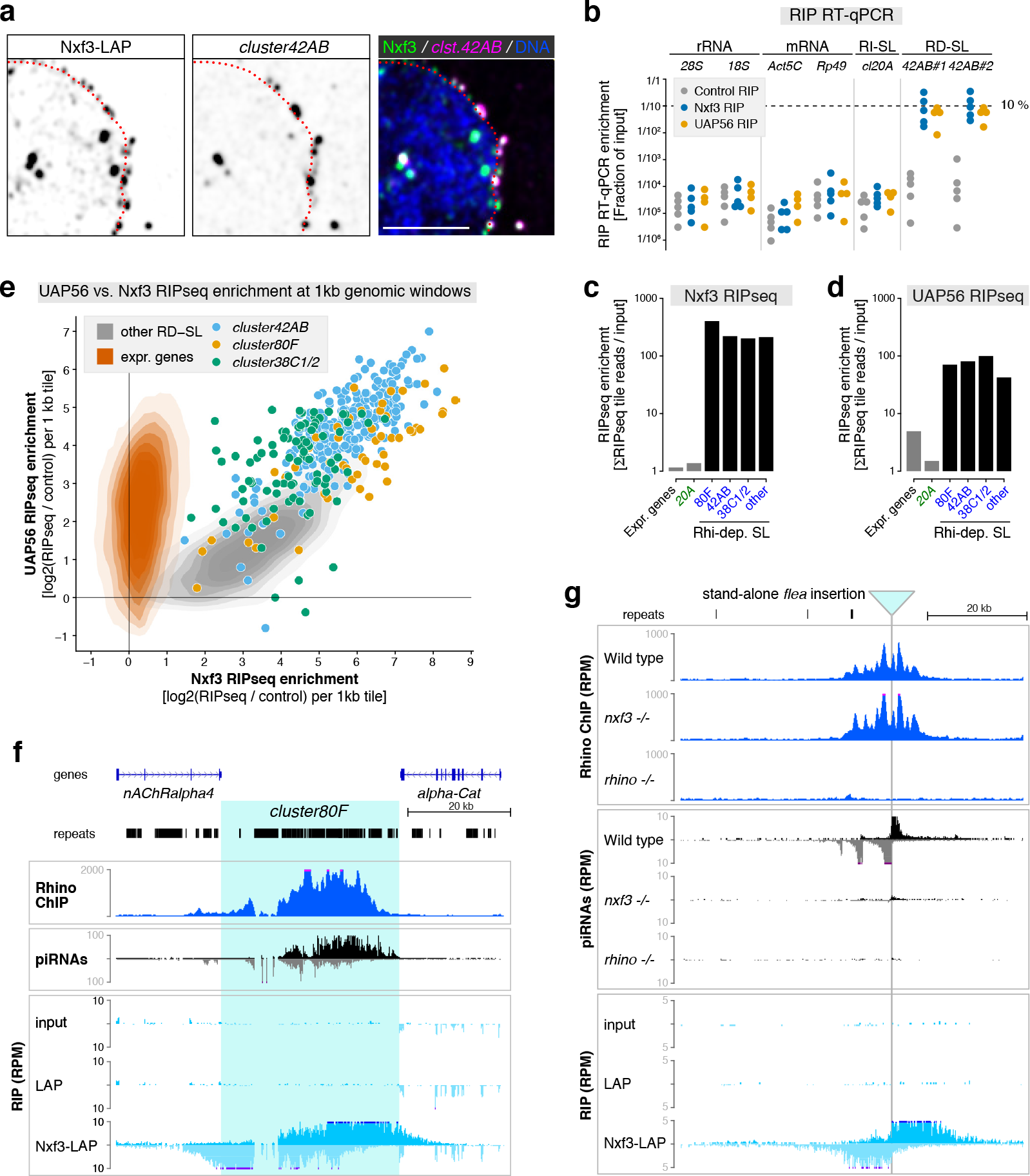
Nxf3 binds piRNA precursors. **a**, Confocal image showing co-localization of Nxf3-LAP with *cluster42AB* transcripts in nucleus and nuage (red dotted line: nuclear envelope; scale bar: 5 µm). **b**, Percent-input values for indicated transcripts present in control-, Nxf3-, or UAP56-RIP eluates determined by quantitative RT-PCR (dots represent independent biological replicates). **c, d**, Fold-enrichment values for indicated transcripts in Nxf3-(**b**) or UAP56-(**c**) RIP eluates relative to input (based on uniquely mapping reads from RIP-seq libraries). **e**, Scatter plot showing enrichments (relative to input) of Nxf3-versus UAP56 RIP-seq reads mapping to 1kb tiles overlapping expressed gene loci (orange; n > 25.000) or to 1kb tiles overlapping Rhino-dependent piRNA source loci (grey; n=2996; 1kb tiles mapping to major piRNA clusters are colored). **f, g** UCSC genome browser panels showing Rhino ChIP-seq reads, piRNAs (reads per million miRNAs), and RIP-seq reads from control (LAP) and Nxf3-LAP eluates mapping to *cluster80F*) (**f**) or a stand-alone *flea* transposon insertion (**g**) (not present in reference genome).

To identify Nxf3 cargo genome-wide, we sequenced libraries generated from the RIP eluates. Transcripts from all major Rhino-dependent piRNA source loci, but not those from Rhino-independent source loci (e.g. *cluster20A*) or mRNAs, were enriched several hundred-fold above background in Nxf3-LAP eluates (Figure 3c). The RNA helicase UAP56 also associated with Rhino-dependent piRNA precursors (Figure 3d) (Zhang et al., 2012; Zhang et al., 2018). Unlike Nxf3, however, UAP56 bound also mRNAs, consistent with its role in mRNA export upstream of Nxf1-Nxt1 (Kohler and Hurt, 2007). While most mRNAs were enriched in the UAP56-RIP libraries, this varied from no measurable enrichment (often for highly abundant transcripts like ribosomal protein-encoding mRNAs) to up to 50-fold enrichment (e.g. transcription factor encoding genes like *vrille* or *schnurri*; Fig. S4c), probably reflecting the ratio of nuclear versus cytoplasmic transcripts for the different mRNAs.

Besides the few major germline piRNA clusters, Rhino is also enriched at hundreds of other genomic loci, mainly located in peri-centromeric heterochromatin (Mohn et al., 2014; Zhang et al., 2014). The RIP libraries revealed that, genome-wide, Rhino-bound loci (a total of 2996 genomic 1kb tiles; RD-SL tiles in Figure 3e) give rise to transcripts that associate with both, Nxf3 and UAP56 (Figure 3e; Figure S4d, e). In contrast, transcripts from tiles overlapping expressed gene loci (~25,000 1kb tiles; not bound by Rhino), were bound by UAP56, but not by Nxf3 (Figure 3e; Figure S4d, e). Our data indicate that, in addition to its role in generating mRNP complexes competent for Nxf1-Nxt1-mediated export, UAP56 may play an analogous function at piRNA clusters in connection with Nxf3-Nxt1.

Inspection of Nxf3-bound transcripts originating from distinct piRNA clusters revealed that top and bottom strand precursors were asymmetrically distributed relative to Rhino’s chromatin occupancy. For example, the first third of *cluster80F* was enriched for bottom strand precursors, while the last third was enriched for top strand precursors (Figure 3f). This pattern is consistent with a model, where high Rhino levels within the cluster drive bidirectional transcription initiation through Moonshiner, and divergent transcription extends for several kilobases (Andersen et al., 2017). At stand-alone transposon insertions, which often act as Rhino-dependent mini-clusters (Mohn et al., 2014; Shpiz et al., 2014), Nxf3 similarly associated with divergent transcripts that bleed into flanking genomic regions from the transposon insertion site (Figure 3g). Rhino occupancy at such transposon insertions was unchanged in *nxf3* mutants, yet piRNA production (as inferred from piRNAs mapping uniquely to the transposon-flanking regions) was lost. We conclude that Nxf3 associates specifically with transcripts originating from Rhino-bound loci genome-wide.

### CG13741/Bootlegger recruits Nxf3 to piRNA clusters

To uncover the molecular basis of Nxf3’s specificity for Rhino-dependent piRNA precursors, we immuno-purified Nxf3-LAP from ovary lysates and identified co-eluting proteins via quantitative mass spectrometry. Besides Nxt1, two additional factors, UAP56 and the uncharacterized protein CG13741 were highly enriched in Nxf3 immuno-precipitates (Figure 4a). In *uap56[28/sz15]* mutants, an allelic combination that specifically prevents UAP56 localization to Rhino-dependent clusters but not its function in mRNA export (Meignin and Davis, 2008; Zhang et al., 2012), Nxf3 levels were reduced (Figure 4b), supporting a functional link between these two factors. However, as UAP56 binds Nxf3-dependent piRNA precursors as well as Nxf1-dependent mRNAs (Figure 3e), it is unlikely to determine Nxf3 specificity. In support of this, *uap56[28/sz15]* mutant germline cells still displayed co-localization of the remaining Nxf3 with Rhino in discrete nuclear foci (Figure 4c).

**Figure 4.**
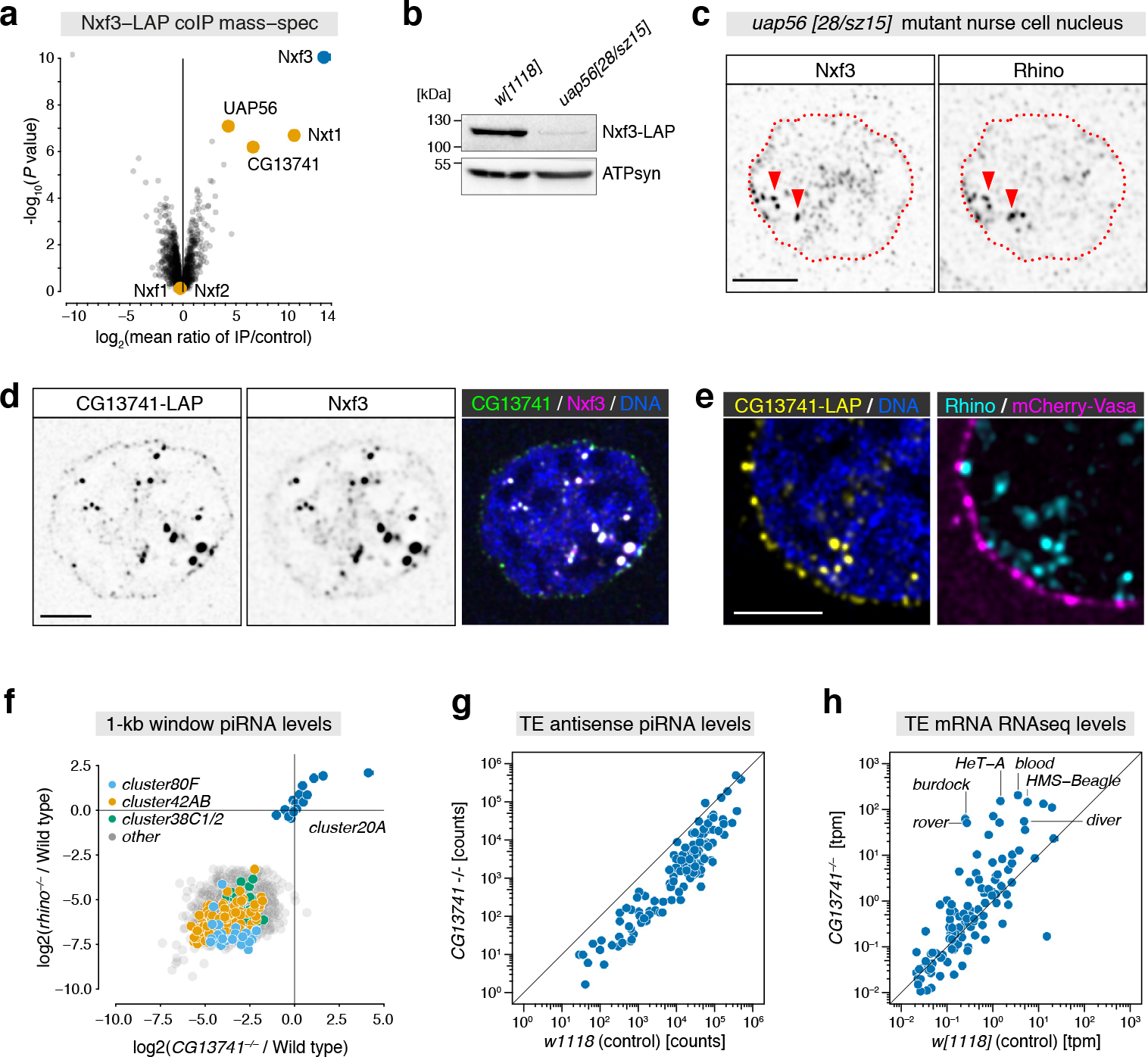
CG13741/Bootlegger interacts with Nxf3. **a**, Volcano plot showing enrichment values and corresponding significance levels for proteins co-purifying with Nxf3-LAP versus control (nuclear LAP) from ovary lysates (n = 4). **b**, Western blot indicating Nxf3-LAP levels in control versus *uap56* mutant ovary lysates (ATP-synthase serves as loading control). **c**, Nxf3-LAP and Rhino localization in nurse cell nucleus of a *uap56* mutant ovary (scale bar: 5 µm; red dotted line: nuclear envelope). **d**, Confocal images showing a nurse cell nucleus (scale bar: 5 µm) indicating co-localization of CG13741-LAP and Nxf3 (antibody stained; we note that Nxf3 in nuage foci is less efficiently detected using antibodies compared to the signal observed from Nxf3-LAP). **e**, Part of a nurse cell nucleus indicating localization of CG13741-LAP, Rhino, and mCherry-Vasa (scale bar: 5 µm). **f**, Scatter plot displaying log2 fold changes of piRNAs mapping uniquely to 1kb tiles from Rhino-dependent piRNA clusters in *rhino* versus *CG13741* mutants (relative to control). Tiles from major piRNA clusters are colored, *cluster20A* tiles serve as Rhino-independent control group. **g, h**, Scatter plots displaying levels of antisense piRNAs (**g**) or levels of sense mRNA reads (**h**) mapping to consensus transposon sequences from control versus *CG13741* mutant ovary samples.

The other Nxf3 interactor, CG13741, was previously identified as a putative piRNA pathway factor in a genetic transposon de-repression screen (Czech et al., 2013). CG13741 lacks recognizable protein domains and is preferentially expressed in ovaries (Brown et al., 2014) (Figure S5a). Using CRISPR-Cas9, we inserted a LAP tag at the C-terminus of the endogenous *CG13741* locus. Imaging of ovaries from CG13741-LAP flies revealed that CG13741 exhibits a dual subcellular localization indistinguishable from that of Nxf3 (Figure 4d), with the nuclear signal corresponding to piRNA clusters (Rhino-positive), and the cytoplasmic signal corresponding to nuage (Vasa-positive; Figure 4e).

To test whether CG13741 is important for Rhino-dependent piRNA clusters and Nxf3 function, we generated *CG13741* frameshift null alleles (Figure S5b, c). Like *rhino* and *nxf3* mutants, *CG13741* mutants were viable yet female sterile, and exhibited loss of piRNAs specifically from Rhino-dependent piRNA source loci (Figure 4f). Consequently, antisense piRNAs targeting germline transposons were reduced (Figure 4g, Figure S5d), and several transposons (e.g. *HMS-Beagle*, *blood*, *HeT-A*, *burdock*) were strongly de-silenced, while mRNA expression was globally unperturbed (Figure 4h; Figure S5e). These findings defined CG13741 as piRNA pathway-specific factor with an essential function at Rhino-dependent piRNA clusters in conjunction with Nxf3. We next determined the genetic dependency between CG13741 and Nxf3. In *CG13741* mutants, Nxf3 protein levels were reduced and the remaining Nxf3 protein showed no enrichment at nuclear Rhino-foci, which remained prominent throughout oogenesis in the absence of CG13741 (Figure 5a, b). These data pointed to a recruitment hierarchy of Nxf3-associated proteins to Rhino-dependent piRNA source loci. We therefore systematically probed the localization of Nxf3, CG13741, Nxt1, Rhino, Cutoff, and Deadlock in ovaries depleted of factors implicated in Rhino biology. Loss of Rhino, Deadlock, or Cutoff (three interdependent proteins co-localizing at nuclear Rhino foci) resulted in reduced levels of Nxf3 and CG13741 proteins (Figure S5f, g), and in their dispersal in nuclei (Figure 5c). Loss of CG13741 had similar effects on Nxf3, but Rhino, Deadlock, and Cutoff still accumulated together in nuclear foci (Figure 5c, Figure S5h). Loss of Nxf3 instead had no impact on CG13741 localization to Rhino/Deadlock/Cutoff foci in the nucleus (Figure 5d; Figure S5h, i). It did, however, prevent the accumulation of Nxt1 at Rhino foci (Figure 5e). Together, these data placed CG13741 downstream of Rhino/Deadlock/Cutoff and upstream of the Nxf3-Nxt1 heterodimer in the nuclear recruitment hierarchy to piRNA clusters. Due to its central role in nuclear export of piRNA precursors—products of Moonshiner-mediated transcription within heterochromatin—we named CG13741 Bootlegger.

**Figure 5.**
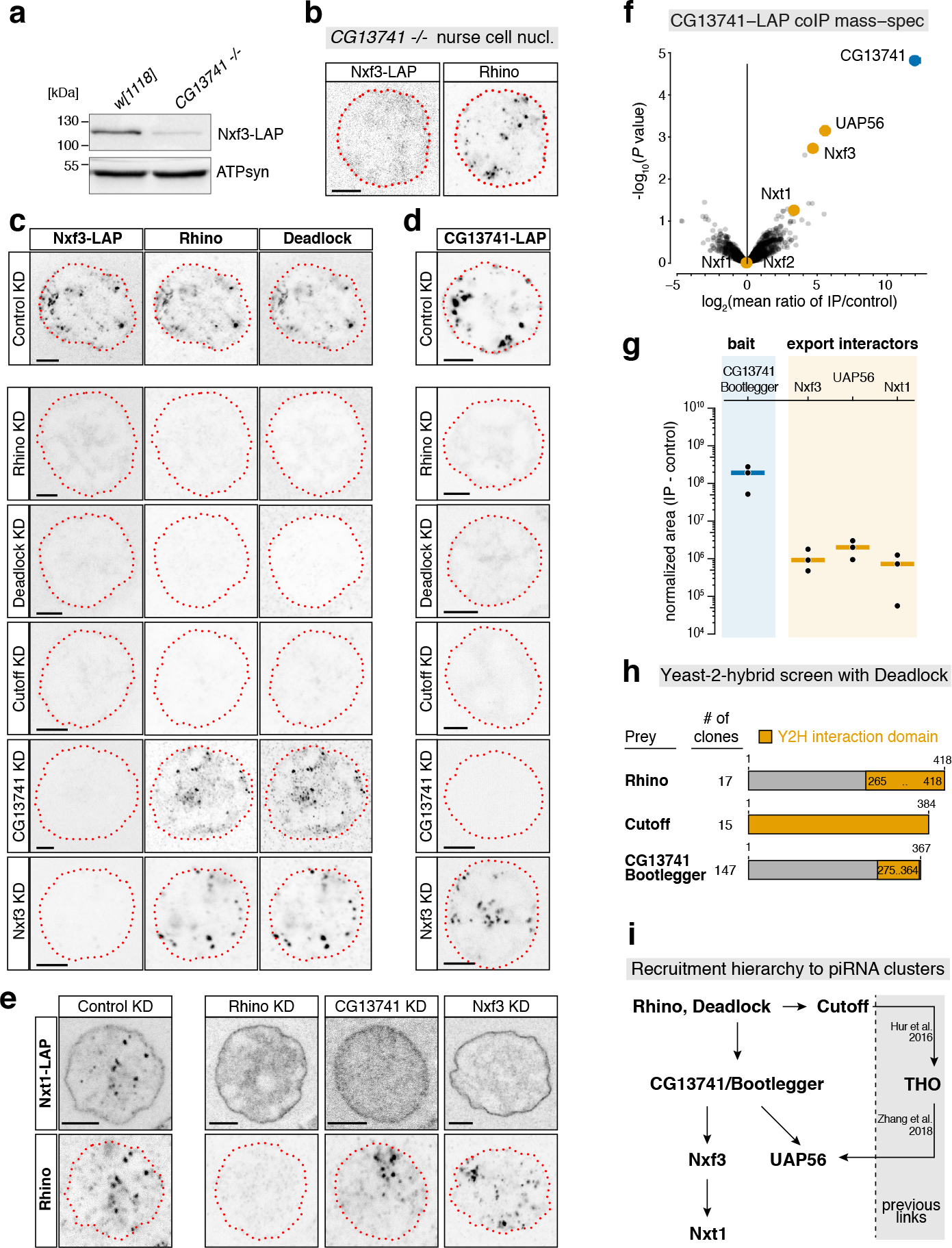
CG13741/Bootlegger recruits Nxf3 to piRNA clusters. **a**, Western blot indicating Nxf3-LAP levels in control versus *CG13741* mutant ovary lysates (ATP-synthase serves as loading control). **b**, Nxf3-LAP and Rhino localization in nurse cell nucleus of a *CG13741* mutant ovary (scale bar: 5 µm; red dotted line: nuclear envelope). **c**, **d**, Confocal images of representative nurse cell nuclei expressing Nxf3-LAP (**c**) or CG13741-LAP (**d**) and sh-lines (KD) targeting indicated genes (red lines: nuclear envelope). **e**, Confocal images of representative nurse cell nuclei expressing Nxt1-LAP and sh-lines (KD) targeting indicated genes (red lines: nuclear envelope). **f**, Volcano plot showing enrichment values and corresponding significance levels for proteins co-purifying with CG13741-LAP versus control (nuclear LAP) from ovaries (n = 3). **g**, Peptide peak intensities (experiment minus respective replicate control) for the CG13741-LAP bait and the proteins indicated in (**d**) (bars: median values). **h**, Cartoon depicting all prey clones retrieved from a yeast two-hybrid screen using full-length Deadlock as bait and employing a *Drosophila* ovarian cDNA library. **i**, Nuclear recruitment hierarchy of indicated proteins to piRNA clusters, including links from the literature.

To understand how Bootlegger connects to the upstream RDC complex at piRNA clusters, we immuno-purified Bootlegger-LAP from ovary lysates and determined co-eluting proteins with quantitative mass-spectrometry. While this confirmed the interaction between Bootlegger, UAP56, and the Nxf3-Nxt1 heterodimer (Figure 5f, g), it did not reveal any link to Rhino/Deadlock/Cutoff. A complementary attempt to identify Bootlegger interactors through a yeast two-hybrid (Y2H) screen failed due to the strong auto-activating character of Bootlegger as bait. We therefore performed a Y2H screen with the Rhino-interactor Deadlock, which acts as adaptor protein by recruiting Cutoff and Moonshiner, two effector proteins involved in piRNA cluster transcription (Andersen et al., 2017; Mohn et al., 2014). From a *Drosophila* ovarian cDNA library, we recovered more than a dozen clones each for the previously identified Deadlock-interactors Rhino and Cutoff (Figure 5h). Notably, more than one hundred sequenced clones represented Bootlegger fragments, all of which shared a ~100 amino acid peptide at the C-terminus. This established a direct molecular link between Bootlegger and Deadlock, and thereby the Rhino/Deadlock/Cutoff complex that defines heterochromatic piRNA clusters.

Nxf3 as well as Bootlegger associate with the general mRNA export factor UAP56 (Figure 4a, 5f). We therefore extended our genetic hierarchy experiments and asked if Nxf3 or Bootlegger also contribute to the recruitment of UAP56 to Rhino-dependent piRNA clusters. Depleting Bootlegger, but not Nxf3, led to a loss of UAP56 enrichment at Rhino foci (Figure S5j). Conversely, in *uap56[28/sz15]* mutants, Bootlegger and Nxf3 levels were diminished, yet both proteins still localized to nuclear Rhino foci (Figure 5a, b; Figure S5k, l). Bootlegger is therefore required for UAP56 to localize to piRNA clusters. Together with published data, this suggests that UAP56 recruitment to RDC requires two molecular interactions, one via Bootlegger and one via the THO complex (Zhang et al., 2018), which is recruited to RDC via Cutoff (Hur et al., 2016). Considering that in *uap56[28/sz15]* mutants, the qualitative pattern of Rhino’s chromatin occupancy is changed (Zhang et al., 2018) our findings are in line with the stronger phenotype of *bootlegger* versus *nxf3* mutants in respect to piRNA levels and transposon de-repression (Figures 2d, e versus Figures 4g, h).

Altogether, our data imply that Rhino, via Deadlock, recruits Bootlegger, which in turn recruits the Nxf3-Nxt1 heterodimer and UAP56 to piRNA source loci, thereby specifying the piRNA precursor export pathway (Figure 5i).

### Nxf3 mediates piRNA precursor export via Crm1

Besides their differing RNA cargo specificity, a second intriguing discrepancy between Nxf3 and the mRNA export receptor Nxf1 is their respective fate after NPC passage. Nxf1 is actively stripped off its RNA cargo right after it reaches the cytoplasmic side of the NPC (Kohler and Hurt, 2007; Lund and Guthrie, 2005). This dissociation is thought to prevent re-import of mRNA into the nucleus and thereby confers directionality to mRNA export. Consistent with this, immunofluorescence experiments (Figure 1d) and immuno-gold electron microscopy (Figure S6a) detected Nxf1 enriched at the nuclear envelope, but not in the cytoplasm. In contrast, Nxf3 is enriched together with piRNA precursors in nuage, a cytoplasmic RNA processing granule (Figure 1h, Figure S1g). This suggested a fundamentally different export mechanism for Nxf3 that ensures transport directionality despite Nxf3 remaining bound to its RNA cargo after NPC passage.

To understand how Nxf3 achieves nuclear export, we compared it to Nxf1, which harbors two nucleoporin binding sites that act synergistically to promote NPC translocation (Braun et al., 2002; Fribourg et al., 2001). The first site is formed by Nxf1’s ubiquitin-associated domain (UBA). Nxf3 lacks a recognizable UBA domain and this nucleoporin binding site is therefore absent (Figure 6a; Figure S6b). Nxf1’s second nucleoporin binding site is formed by the NTF2-like domain, which binds to Nxt1. Nxf1-Nxt1 hetero-dimerization via this interaction is required for Nxf1’s nuclear shuttling ability (Katahira et al., 2002; Levesque et al., 2001). We therefore investigated Nxt1’s role in Nxf3 export. Using a heterologous co-immuno-precipitation assay in Schneider cells, we verified that Nxt1 binds Nxf3’s NTF2-like domain (Figure S6c). This interaction was lost upon mutation of Asp-434 (D434), whose equivalent residue in Nxf1 forms a conserved salt bridge to Nxt1 (Figure S6c) (Kerkow et al., 2012). We engineered the D434R allele in the endogenous, LAP-tagged *nxf3* locus of flies. Nxf3[D434R] accumulated at reduced levels *in vivo* and was unable to bind Nxt1 above background (Figure 6b; Figure S6d). However, Nxf3[D434R] still interacted with UAP56 and Bootlegger (Figure 6b), co-localized with Rhino at piRNA clusters in the nucleus (Figure S6e), and was enriched in cytoplasmic nuage albeit to a slightly reduced extent (Figure 6c, Figure S6f). *nxf3[D434R]* mutants were, however, sub-fertile and exhibited de-silencing of transposons (Figure S6g). Consistent with this, nuage-localized *cluster42AB* transcripts and piRNAs from Rhino-dependent source loci were reduced in *nxf3[D434R]* mutants (Figure S6h, i). Besides stabilizing Nxf3, Nxt1 might therefore contribute to RNA cargo binding, similar to what is known for Nxf1-Nxt1 (Aibara et al., 2015; Katahira et al., 2015). Altogether, our data demonstrate that Nxt1 is important for Nxf3 function, and suggest that Nxf3 achieves nuclear export independent of nucleoporin binding sites that are critical for Nxf1.

**Figure 6.**
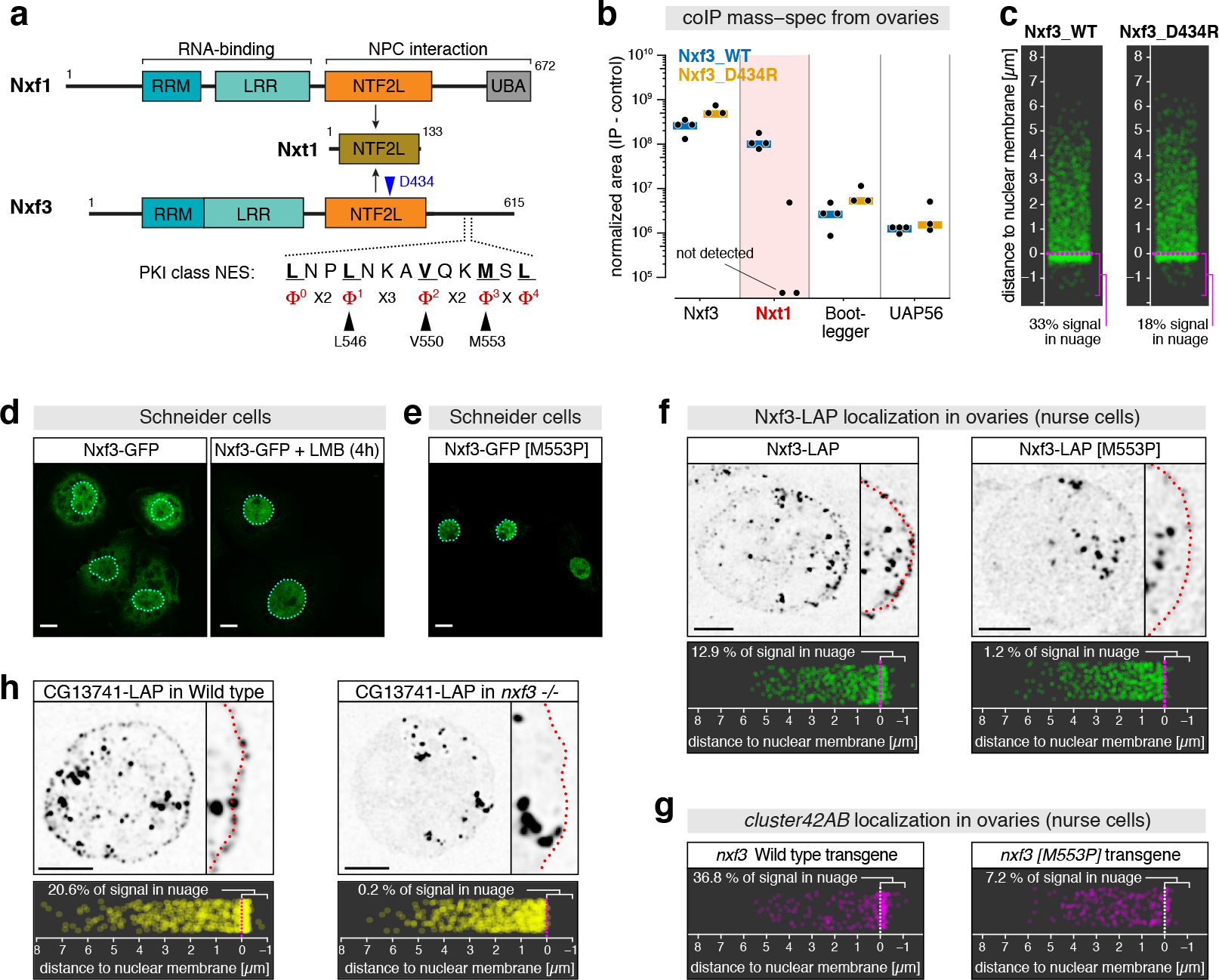
Nxf3 utilizes Crm1 for nuclear export. **a**, Cartoon depicting protein domain composition of Nxf1, Nxf3, and Nxt1 (RRM: RNA binding motif; LRR: Leucine-rich repeat; NTF2-like domain; UBA: Ubiquitin-associated domain). The PKI type NES in Nxf3 is indicated. **b**, Absolute peptide peak intensities for indicated Nxf3 interactors from co-immuno-precipitation of wildtype Nxf3-LAP (WT) and Nxt1-interaction mutant Nxf3-LAP (D434R). **c**, Jitter plots showing positions of Nxf3-LAP foci in nurse cells relative to the nuclear envelope (n = 1500 foci; see Figure S6f for representative confocal image). **d**, Confocal images showing sub-cellular localization of Nxf3-GFP in untreated (left) or LMB-treated (right) S2 cells (scale bar: 5 µm). **e**, Confocal image showing sub-cellular localization of Nxf3-GFP [M553P] (see panel **a**) in S2 cells (scale bar: 5 µm). **f**, Confocal images (scale bar: 5 µm; red dotted line: nuclear envelope) showing localization of Nxf3-LAP (left) or Nxf3-LAP [M553P] (right) in nurse cells (Nxf3-LAP expressed under the *nanos* promoter in *nxf3* mutant background). Jitter plots below indicate the distance of respective protein foci to nuclear envelope (n = 501 foci analyzed per genotype). **g**, Jitter plots showing positions of *cluster42AB* FISH foci in nurse cells of indicated genotype relative to the nuclear envelope (n = 223 foci; see Figure S7e for representative confocal image). **h**, Confocal images of representative nurse cell nuclei (scale bar: 5 µm; magnified panels show part of nucleus; red dotted line: nuclear envelope) showing localization of CG13741/Bootlegger. Jitter plots below indicate the distance of Bootlegger protein foci to the nuclear envelope (n = 572 foci analyzed per genotype).

Human NXF3, an orphan NXF variant more closely related to human NXF1 than to *Drosophila* Nxf3, achieves NPC translocation via a leucine-rich nuclear export signal (NES) that is recognized by Crm1, the principal cellular exportin (Yang et al., 2001). When expressed in S2 cells, *Drosophila* Nxf3 was mostly cytoplasmic and accumulated to only moderate levels in the nucleus (Figure 6d). Upon treating S2 cells with the Crm1 inhibitor Leptomycin B (LMB) (Kudo et al., 1999), exogenous Nxf3 became entirely nuclear (Figure 6d, Figure S7a), as did endogenous Nxf3 in LMB-treated ovaries (Figure S7b). We identified a conserved PKI-type NES candidate downstream of Nxf3’s NTF2-like domain that matches all criteria defined for this NES type (Figure 6a; Figure S6b) (Guttler et al., 2010). Independent point mutations in this NES (construct 1: L546A + V550A; construct 2: M553P) rendered Nxf3 entirely nuclear in S2 cells (Figure 6a, e; Figure S7c), supporting a role for this conserved motif as bona fide NES. We did not succeed in generating flies with NES mutations in the endogenous *nxf3* locus, and therefore complemented *nxf3* mutants with a germline-expressed *nxf3* cDNA construct carrying the M553P mutation. Nxf3[M553P] localized to nuclear Rhino foci (Figure 6f), was expressed at levels comparable to the ectopically expressed wild-type protein (Figure S7d), but did not accumulate efficiently in cytoplasmic nuage (Figure 6f). Similarly, *cluster42AB* transcripts accumulated only weakly in nuage of *nxf3[M553P]* mutant ovaries (Figure 6g; Figure S7e). These findings indicate that Nxf3 utilizes an NES in order to achieve efficient nuclear export via Crm1. *nxf3[M553P]* mutant flies exhibited strong sterility only after ~5 days (Figure S7f). We speculate that export-impaired Nxf3 still stabilizes piRNA precursors in the nucleus, and that due to leakage of precursors to the cytoplasm the NES-mutant exhibits a milder phenotype compared to *nxf3* null mutants.

Our data reveal that Nxf3, rather than using direct nucleoporin-binding sites like Nxf1, has evolved an NES motif in order to tap into the Crm1-dependent nuclear export system. Remarkably, in *nxf3* mutants, where Vasa-positive nuage granules remain abundant (Figure S7g), Bootlegger is entirely nuclear and fails to enrich in nuage (Figure 6h). Thus, by utilizing the Crm1 system, Nxf3 achieves export not only for its RNA cargo but also for Bootlegger. In contrast to the rapid cytoplasmic dissociation of Nxf1 from its cargo mRNAs, Crm1-translocated Nxf3 can remain bound to its RNA cargo and Bootlegger after nuclear exit. We speculate that this allows Nxf3/Bootlegger to direct piRNA precursors to nuage, where PIWI-clade proteins loaded with complementary piRNAs will initiate piRNA biogenesis (Han et al., 2015; Homolka et al., 2015; Mohn et al., 2015; Webster et al., 2015).

## DISCUSSION

In this work, we uncover the Nxf3/Bootlegger pathway, which transports RNA Polymerase II-generated piRNA precursor transcripts emanating from heterochromatin to peri-nuclear nuage where they are processed into piRNAs. This pathway bypasses nuclear RNA surveillance mechanisms that would otherwise degrade the unprocessed precursor RNAs, and enables their nuclear export and delivery to nuage. Retroviruses also must bypass nuclear RNA quality control mechanisms to export their unprocessed genomic RNA. Specific hairpins in the retroviral RNA recruit the mRNA export receptor Nxf1/Tap (e.g. Mason-Pfizer monkey virus) (Gruter et al., 1998), or exploit the exportin Crm1 through a virally encoded adaptor protein (e.g. HIV) (Cullen, 2000). Due to their high sequence diversity, it would be challenging for piRNA precursors to recruit export machinery via sequence or structural features. Instead, we suggest that specificity in piRNA precursor export is ultimately achieved through Rhino, which recruits the specialized NXF variant, Nxf3, to nascent transcripts at heterochromatic piRNA clusters. The Nxf3/Bootlegger pathway highlights Rhino’s central role in facilitating productive expression of transposon sequence information within heterochromatin (Figure 7). Via its adaptor protein Deadlock, Rhino connects H3K9me3 marks at piRNA clusters to a set of effector proteins that allow transcription initiation (through the TFIIA-L homolog Moonshiner), prevent transcription termination (through the Rai1 homolog Cutoff), and direct nuclear export of the emerging unprocessed RNAs (through the Nxf1 homolog Nxf3).

**Figure 7.**
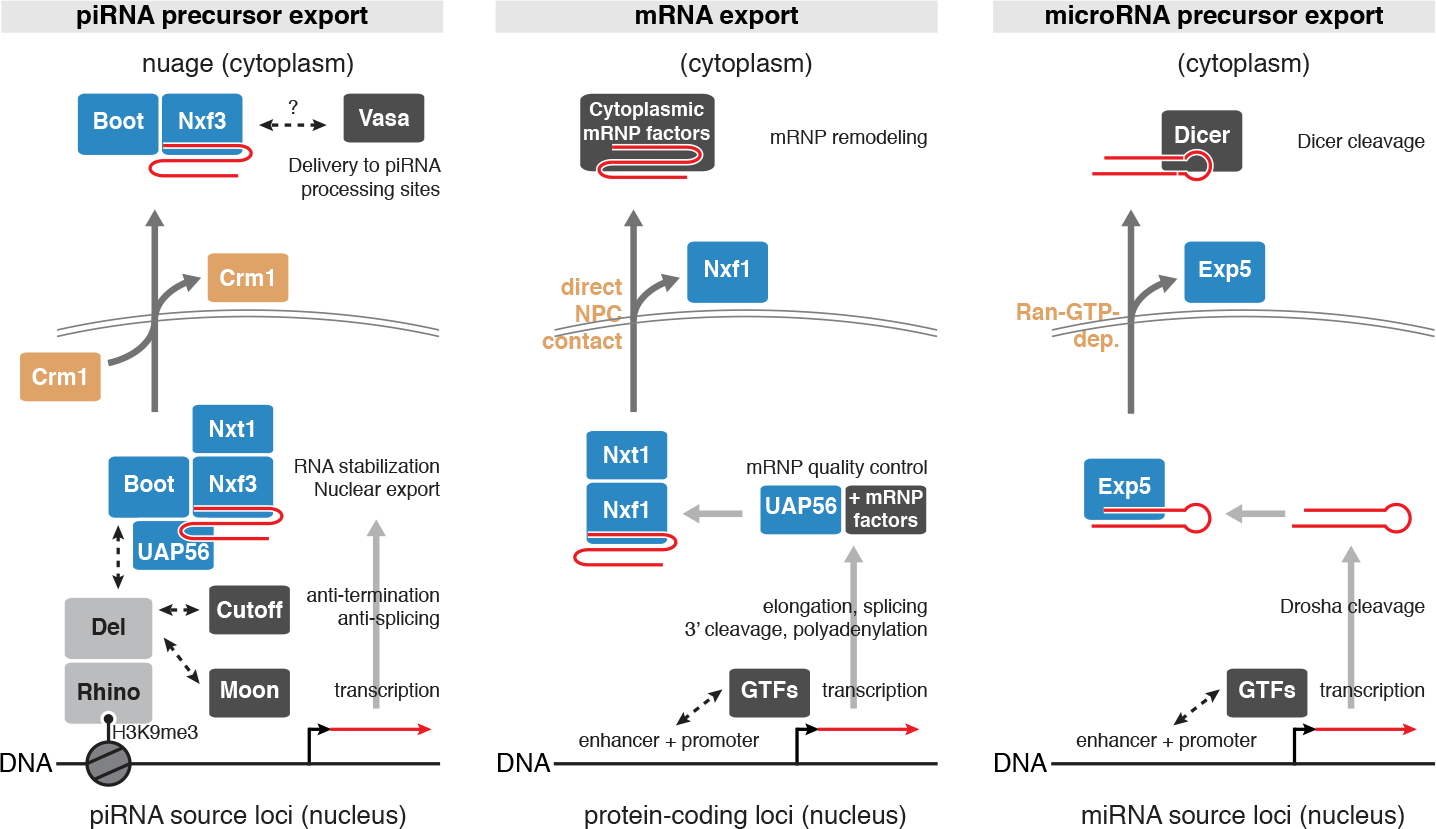
Comparison of nuclear RNA export pathways. Schematics of the Nxf3/Bootlegger pathway in comparison to mRNA export and pre-miRNA export (GTF: general transcription factors; Exp5: Exportin-5).

Nxf3 and Bootlegger co-purify with UAP56, a nuclear DEAD-box ATPase that facilitates loading of the Nxf1-Nxt1 heterodimer onto export-competent mRNA. In the context of mRNA export, UAP56 functions together with the pentameric THO complex (Heath et al., 2016; Ren et al., 2017). Recent work indicates a role for UAP56 and THO in piRNA cluster transcription and in defining Rhino’s chromatin occupancy (Zhang et al., 2018). Based on our findings, we speculate that UAP56 and THO, analogous to their function in mRNA export, are also required to license loading of Nxf3-Nxt1 onto piRNA precursors. Biochemical and structural studies will be crucial to understanding how Nxf3 molecularly diverged from the Nxf1 export receptor to become an export adaptor for Crm1-dependent translocation of transcripts from specific genomic loci, yet still requiring factors in common with Nxf1 like UAP56 and THO (Figure 7).

While the piRNA precursor-specialized Nxf3/Bootlegger pathway evolved from the general mRNA export machinery, it also shares similarities to pre-microRNA export, which is dependent on Exportin-5 (Bohnsack et al., 2004; Lund et al., 2004; Yi et al., 2003) (Figure 7). In both cases, the export machinery not only facilitates passage through the NPC, but binding of the export factors to cargo RNA also protects against nuclear RNA degradation. Similarly, and in contrast to mRNA export, both pathways utilize the Ran-GTP gradient, either directly (Exportin-5) or indirectly via Crm1 (Nxf3). As the Ran-GTP gradient confers directionality to nuclear transport pathways (here via dissociation of Crm1 and Nxf3 in the cytoplasm), we suggest that this allows Nxf3 to remain bound to its RNA cargo after nuclear export. It is striking that human NXF3 has independently evolved an export dependency on Crm1, suggesting that also human NXF3 might hold on to its cargo after NPC translocation. Remaining cargo-bound enables Nxf3 and Bootlegger to escort piRNA precursors to nuage, a biomolecular condensate formed by liquid-liquid phase separation (Nott et al., 2015) where the piRNA biogenesis machinery is concentrated. Future molecular and biophysical investigations of the Nxf3/Bootlegger pathway promise to uncover how piRNA precursors are directed to nuage.

Notably, *Drosophila* Nxf2 is essential for heterochromatin formation downstream of nuclear Piwi (J. Batki, J. Schnabl, J. Brennecke, manuscript in preparation). This points to the NXF protein family as a hot spot for genetic innovation through gene duplication and exaptation. In vertebrates, Nxf1/Tap has also diversified into multiple NXF variants. Mammalian NXF2 and NXF3 are preferentially expressed in gonads, and *Nxf2* mutant mice are male sterile (Pan et al., 2009). We propose that neo-functionalized NXF variants have evolved throughout the animal kingdom to serve key functions in genome defense.

## Supporting information

Supplementary Table 1 Drosophila melanogaster fly strains

Supplementary Table 2 Oligo sequences

Supplementary Table 3 Antibody list

Supplementary Table 4 Stellaris probe sequences

Supplementary Table 5 Antisense rRNA oligo list

Supplementray Table 6 Sequence accessions

## ACKNOWLEDGMENTS

We thank D. Handler for bioinformatics help, P. Duchek and J. Gokcezade for generating CRISPR-edited and transgenic flies, K. Mechtler and his team for mass spectrometry, M. Novatchkova for sequence and phylogenetic analysis, the Vienna Biocenter Core Facilities (VBCF) Next Generation Sequencing unit for deep sequencing, VBCF Protein Technologies Facility for recombinant protein expression, the VBCF EM Facility for electron microscopy and the BioOptics Facility, in particulay P. Pasierbek, for their support. Bill Theurkauf, Ilan Davis and Elizabeth Gavis provided fly lines. We thank Torbein Heick Jensen, Clemens Plaschka, Ulrich Hohmann, and Guy Riddihough from Life Science Editors (http://lifescienceeditors.com) for comments on the manuscript. The Brennecke lab is supported by the Austrian Academy of Sciences, the European Research Council (ERC-2015-CoG - 682181), and the Austrian Science Fund (F 4303). P.R.A. is supported by the Novo Nordisk Foundation (NNF14OC0009189 and NNF17OC0030954), and M.E. by a DOC Fellowship from the Austrian Academy of Sciences.

## AUTHOR CONTRIBUTIONS

M.E., L.T., F.P. and P.R.A. performed all molecular biology and fly experiments. K.M. established LAP-tagged fly lines, T. L. and M.E. performed the image analysis for sub-cellular protein localization. M.E., J.B. and P.R.A. performed the computational analyses. P.R.A. and J.B. supervised the project. M.E., P.R.A. and J.B. wrote the paper.

## MATERIALS & METHODS

### Fly Husbandry

Flies were kept at 25°C under light/dark and humidity (60%) control. For ovary dissection, 3-5 days old flies were kept in cages with yeast supply on apple juice plates for two days and then dissected after brief immobilization by CO_2_ anesthesia. A list of fly strains including genotypes, identifiers and original sources can be found in Supplementary Table 1. The listed stocks are available via VDRC (http://stockcenter.vdrc.at/control/main).

### Generation of transgenic fly strains

We generated fly strains harboring short hairpin RNA (shRNA) expression cassettes for germline knockdown by cloning shRNA sequences into the Valium-20 vector (Ni et al., 2011) modified with a *white* selection marker (oligos are listed in Supplementary Table 2). Tagged fly stains were generated via insertion of desired tag sequences into locus-containing Pacman clones via bacterial recombineering (Ejsmont et al., 2011; Venken et al., 2009). Wild type and NES-mutant Nxf3 transgenic flies were generated by inserting germline expression constructs (*nanos* promoter and *vasa* 3’ UTR) plus the corresponding coding sequences into the attP40 landing site (FlyBase ID: FBti0114379). Of note, the annotated coding sequence of Nxf3 (FlyBase ID: FBgn0263232) lacks the first 56 amino acids of the Nxf3 protein. Using CAP-seq data, we identified a transcription start site (TSS) upstream of the annotated TSS. Germline expression of the currently annotated Nxf3 tagged at the C-terminus with a LAP-tag gives rise to an aberrant localization pattern. However, expression of the full ORF gives a localization pattern that is identical to the endogenously engineering locus, both of which are functional rescues. Furthermore, the non-annotated part upstream of the annotated TSS is conserved in other fly species.

### Generation of mutant fly strains

Mutant alleles for *nxf3*, *CG13741*, *uap56 [E245K]*, and *nxf3[D434R]* were generated as described in (Gokcezade et al., 2014) through injection of pDCC6 plasmids modified to express guide-RNAs, along with ssDNA repair templates in case of point mutant generation. Oligos and targeting sequences are given in Supplementary Table 2.

### Generation of endogenous knock-in fly strains

CRISPR-Cas9-mediated generation of endogenously tagged lines was through injection of pU6-BbsI-chiRNA plasmids (Addgene:466294; (Gratz et al., 2013)) along with pBS donor plasmids containing 1 kb long homology arms into act-Cas9 embryos (Bloomington stock: 54590; (Port et al., 2014)). Oligos and targeting sequences are listed in Supplementary Table 2.

### *Drosophila* Schneider 2 (S2) cell culture

*Drosophila* Schneider 2 (S2) cells (in-house stock regularly tested to be virus- and mycoplasma-free) were grown at 25°C in S2 cell media supplemented with 10% fetal bovine serum (Thermo Fisher Scientific).

### Protein co-immunoprecipitation from S2 cell lysates

4 million S2 cells were transfected with 2 µg DNA using Cell Line Nucleofector kit V (Amaxa Biosystems) with the program G-030. S2 cells were co-transfected with plasmids expressing FLAG-tagged and GFP-tagged proteins driven by the actin5C promotor. After 48h of incubation at 25°C, S2 cells were collected through centrifugation at 600g. The pellet was resuspended in 100 µl lysis buffer (30 mM HEPES pH 7.4, 150 mM NaCl, 2 mM MgCl_2_, Triton X100 0.2%), freshly supplemented with 1 mM DTT, 1 mM Pefabloc and Complete Protease Inhibitor Cocktail (Roche). Lysis was carried out at 4°C for 20 min followed by 10 min centrifugation at 18,000g at 4°C. Protein concentration was measured using standard Bradford assay and 500 µg of total protein were incubated with GFP-Trap magnetic beads (ChromoTek) for 3h at 4°C. Beads were washed 3× 10min with lysis buffer and proteins were eluted in 1× SDS loading buffer at 95°C.

### Western blot analysis

Western blotting was done following standard protocols. Samples were resolved by SDS-polyacrylamide gel electrophoresis and transferred to a 0.45 µm (Bio-Rad) nitrocellulose membrane. The membrane was blocked in 5% skimmed milk powder in PBS supplemented with 0.01% TritonX100 (PBX) for 30 min and incubated with primary antibody overnight at 4°C. After three washes with PBX, the membrane was incubated with HRP-conjugated secondary antibody for 2h at room temperature, followed by three washes with PBX. Subsequently, the membrane was covered with Clarity Western ECL Blotting Substrate (Bio-Rad) and imaged using the ChemiDoc MP imaging system (Bio-Rad). All relevant antibodies are listed in Supplementary Table 3.

### Recombinant protein expression in insect cells

HIS-GST-Nxf3 (66-336) was cloned into pGB vector. The resulting plasmid was transposed into the EmBacY bacmid backbone and transfected into *Spodoptera frugiperda Sf9* cells to generate a baculovirus expressing the construct. The virus was used to infect *Trichoplusia ni High5* cells and expression was performed at 21°C. Cells were harvested 4 days after growth arrest.

### His affinity purification

*High5* cells were lysed in lysis buffer (LB) (50 mM Tris-HCL pH=8, 150 mM NaCl, 10% glycerol, 10 mM Imidazole) freshly supplemented with 1 mM Pefabloc, 5 mM ß-mercapto-ethanol and Benzonase (~10U/ml) for 30 min at 4°C. The lysate was cleared by centrifugation. Purification was done using an AKTA Purifier FPLC system on a HisTrap FF 5 ml column (GE Healthcare). The column was washed and equilibrated with 10 volumes of LB prior to sample loading. Bound protein was eluted in LB supplemented with 250 mM Imidazole and subsequently analyzed by SDS-PAGE and InstantBlue (Expedeon) staining.

### Antibody generation

Mouse monoclonal antibodies against His-tagged Nxf3 (amino acids T336-L614), His-tagged CG13741 (full length), and Deadlock (amino acids P470-P669) were generated by the Max F. Perutz Laboratories Antibody Facility. Nxf3 and CG13741 antigens were cloned in pET-15b and transformed in BL21(DE3) *E. coli*, and purified using Ni-NTA resin (Qiagen) according to standard protocols. The deadlock antigen was cloned downstream of a TEV recognition site in pMAL-c2x construct to produce a maltose-binding protein (MBP)-tagged peptide, which was purified using amylose resin (NEB). The MBP-tag was cleaved by TEV protease treatment.

### Electrophoretic mobility shift assay (EMSA)

2.5 nmol [32P] 5’-labelled single stranded 35nt RNA (CUCAUCUUGGUCGUACGC GGAAUAGUUUAAACUGU) was equilibrated with various concentrations of recombinant 6xHis-GST-Nxf3 (66-336) or GST in 10 µl total volume in EMSA binding buffer (10 mM Tris-HCL pH=8.5, 1 mM EDTA, 100 mM KCl, 5% glycerol), freshly supplemented with RNase inhibitor, 1 mM DTT and BSA (10 µg/ml). The reactions were incubated at 4°C for 20 min. 2 µl of EMSA loading buffer (50% glycerol, 0.075% bromophenol blue) was added and the total reaction volumes were analyzed on a pre-run 4.8% PAGE gel in 0.5 × TBE. The radioactive bands were visualized with an Amersham Typhoon Imaging System (GE Healthcare).

### Protein co-immunoprecipitation from ovary lysates

Ovary samples (100 to 200 ovary pairs) were dissected and transferred to ice-cold PBS. After PBS removal, ovaries were then snap-frozen for later processing. For immunoprecipitation, ovary samples were homogenized with 20-30 strokes of a glass douncer with a tight pestle in 1 mL Ovary Protein Lysis Buffer (20 mM Tris-HCl pH 7.5, 150 mM NaCl, 2 mM MgCl_2_, 10 % glycerol, 1 mM DTT, 1 mM PefaBloc, 0.2 % NP-40). Homogenates were transferred to 1.5 mL low-retention tubes and incubated on ice for 15 minutes with occasional inversion. The lysates were cleared by centrifugation for 5 minutes at 16.000*g*. Cleared lysates were added to 20 µL anti-FLAG M2 magnetic bead (Sigma) solution (50% beads per total volume with Beads Buffer (20 mM Tris-HCl pH 7.5, 150 mM NaCl)). Lysate-bead mixtures were incubated 3 hours at 4°C with rotation and then washed four times 10 minutes in Ovary Protein Lysis Buffer followed by six quick rinses in Co-IP Wash Buffer (20 mM HEPES pH 7.4, 150 mM NaCl, 2 mM MgCl_2_). Pelleted magnetic beads were stored at 4°C in 10 µL Co-IP Wash Buffer until processing for mass spectrometry analysis the following day.

### Mass Spectrometry Analyses

Co-immuno-precipitated proteins were subjected to on-bead digestion with LysC. The hereby eluted peptides were digested with Trypsin overnight. The resulting peptides were analyzed using a Dionex UltiMate 3000 HPLC RSLC nano system coupled to a Q Exactive mass spectrometer equipped with a Proxeon nanospray source (Thermo Fisher Scientific). Peptides were eluted using a flow rate of 230 nl min^−1^, and a binary 3h gradient, respectively 225 min and the data were acquired with the mass spectrometer operated in data-dependent mode with MS/MS scans of the 10 most abundant ions. For peptide identification, the RAW-files were loaded into Proteome Discoverer (version 2.1.0.81, Thermo Scientific) and the created MS/MS spectra were searched using MSAmanda v1.0.0.6186 (Dorfer et al., 2014) against *Drosophila melanogaster* reference translations retrieved from Flybase (dmel_all-translation-r6.17). An in-house-developed tool apQuant (Doblmann et al., 2018) was used for the peptide and protein quantification (IMP/IMBA/GMI Protein Chemistry Facility; http://ms.imp.ac.at/index.php?action=apQuant). Average enrichment between bait and control IP experiments were plotted against p values calculated using the limma R package (Smyth, 2004).

### RT-qPCR analysis of transposon expression

Five pairs of dissected ovaries were homogenized in TRIzol reagent followed by RNA purification according to the manufacturer’s protocol. 1 µg of total RNA was digested with RQ1 RNase-free DNase (Promega) and then reverse transcribed using anchored oligo dT primers and Superscript II (Invitrogen) following standard protocols. cDNA was used as template for RT-qPCR quantification of transposon mRNA abundances (for primer sequences see Supplementary Table 2).

### Immuno-gold Electron Microscopy

Freshly dissected ovaries were fixed in 2% paraformaldehyde and 0.2% glutaraldehyde (both EM-grade, EMS, USA) in 0.1 M PHEM buffer (pH 7) for 2h at RT, then overnight at 4°C. The fixed ovaries were embedded in 12% gelatin and cut into 1 mm^3^ blocks which were immersed in 2.3 M sucrose overnight at 4°C. These blocks were mounted onto Leica specimen carrier (Leica Microsystems, Austria) and frozen in liquid nitrogen. With a Leica UCT/FCS cryo-ultra-microtome (Leica Microsystems, Austria) the frozen blocks were cut into ultra-thin sections at a nominal thickness of 60nm at −120°C. A mixture of 2% methylcellulose (25 centipoises) and 2.3 M sucrose in a ratio of 1:1 was used as a pick-up solution. Sections were picked up onto 200 mesh Ni grids (Gilder Grids, UK) with a carbon coated formvar film (Agar Scientific, UK). Fixation, embedding and cryo-sectioning as described by (Tokuyasu, 1973). Prior to immuno-labeling, grids were placed on plates with solidified 2% gelatin and warmed up to 37 °C for 20 min to remove the pick-up solution. After quenching of free aldehyde-groups with glycin (0.1% for 15 min), a blocking step with 1% BSA (fraction V) in 0.1 M Sörensen phosphate buffer (pH 7.4) was performed for 30 min. The grids were incubated in primary antibody (rabbit polyclonal against GFP; ab6556, Abcam, UK), diluted 1:125 in 0.1 M Sörensen phosphate buffer over night at 4°C, followed by a 2h incubation in the secondary antibody (goat-anti-rabbit antibody coupled with 6 nm gold; GAR 6 nm, Aurion, The Netherlands), diluted 1:20 in 0.1 M Sörensen phosphate buffer, performed at RT. The sections were stained with 4% uranyl acetate (Merck, Germany) and 2% methylcellulose in a ratio of 1:9 (on ice). All labeling steps were done in a wet chamber. The sections were inspected using a FEI Morgagni 268D TEM (FEI, The Netherlands) operated at 80kV. Electron micrographs were acquired using an 11 megapixel Morada CCD camera from Olympus-SIS (Germany).

### Immunofluorescence staining of ovaries

5-10 ovary pairs were dissected into ice-cold PBS and subsequently fixed by incubation for 20 minutes at room temperature in IF Fixing Buffer (4 % paraformaldehyde, 0.3 % Triton X-100, 1× PBS) with rotation. Fixed ovaries were washed three times 10 minutes in PBX (0.3 % Triton X-100, 1× PBS) and blocked with BBX (0.1% BSA, 0.3 % Triton X-100, 1× PBS) for 30 minutes, all at room temperature with rotation. Primary antibody incubation was performed by overnight incubation at 4°C with antibodies diluted in BBX followed by three 10 minute-washes in PBX. Ovaries were then incubated overnight at 4°C with fluorophore-coupled secondary antibodies, washed once in PBX, incubated with Alex Flour-conjugated wheat germ agglutinin (Thermo Fisher Scientific) at a 1:200 dilution in PBX for 20 minutes, and washed three times in PBX with the second wash done with DAPI added to the PBX to stain DNA (1:50.000 dilution). The final wash buffer was carefully removed, and each sample was resuspended in ~40 µl Prolong Diamond mounting medium and 10 µl of 0.5µm TetraSpeck beads (Thermo Fisher Scientific). The samples were imaged on a Zeiss LSM-880 Axio Imager confocal-microscope and the resulting images processed using FIJI/ImageJ (Schindelin et al., 2012). TetraSpeck beads were imaged separately in a Z-stack of 320 nm. All relevant antibodies are listed in Supplementary Table 3.

### Immunofluorescence staining of S2 cells

24h after transfection, 30 000 cells were seeded onto removable chambered cover glass (Grace Bio-Labs), freshly coated with Concanavalin A, for 3h. Cells were washed with PBS for 5 min and subsequently fixed with formaldehyde solution (4% formaldehyde in PBS) for 20 min at room temperature. If indicated, cells were treated with 20 ng/ml Leptomycin B (LMB, Sigma Aldrich) prior to fixation. Fixed cells were washed twice with PBS for 5 min, permeabilized with 0.3% TritonX100 in PBS for 10 min and washed again with PBS for 5 min. Blocking was done in BBX (PBX (0.01% TritonX100 in PBS) + 1% BSA) for 20 min. Cells were incubated with primary antibody in BBX at 4°C overnight. Following three washing steps with PBX, cells were incubated with fluorophore-conjugated secondary antibody in BBX for 2h at room temperature in the dark. Stained cells were washed three times with PBX. DAPI stain (1: 5000) was included in the second wash. The coverglass was removed from culture chambers and each sample was mounted using one drop of ProLong Diamond Antifade Mountant (Thermo Fisher Scientific). Mounted samples were imaged with a Zeiss LSM-880 confocal microscope and the images were processed using FIJI/ImageJ. All relevant antibodies are listed in Supplementary Table 3.

### RNA Fluorescent In Situ Hybridization (FISH)

Freshly dissected ovary pairs (5-10 for each sample) were fixed in formaldehyde solution (4% formaldehyde, 0.15% Triton X-100 in PBS) for 20 minutes at room temperature, washed three times for 10 minutes in 0.3% Triton X-100/PBS, and permeabilized overnight in 70% ethanol at 4 °C. Permeabilized ovaries were rehydrated for 5 minutes in RNA FISH wash buffer (10% (v/w) formamide in 2× SSC). Ovaries were resuspended in 50 µl hybridization buffer (10% (v/w) dextran sulfate and 10% (v/w) formamide in 2× SSC) containing 0.5 µl of each 25 µM RNA FISH probe set solution (Stellaris; see Supplementary Table 4 for probe sequences) and hybridized overnight at 37 °C with rotation. Next, ovaries were rinsed twice in RNA FISH wash buffer, resuspended in RNA FISH wash buffer containing wheat germ agglutinin-coupled Alexa Fluor 488 or 647 conjugate (WGA-488) at a final concentration of 5 ng/µl and then incubated for 1h at room temperature with agitation. Ovaries were then subjected to four washes: once for 30 minutes at room temperature in RNA FISH wash buffer, once for 10 minutes in a DAPI/2xSSC solution and finally twice for 10 min in 2× SSC buffer. The final wash buffer was carefully removed, and each sample was resuspended in ~40 µl Prolong Diamond mounting medium and mounted on a microscopy slide. The mounted samples were allowed to equilibrate for at least 24 h before imaging on a Zeiss LSM 880 confocal microscope equipped with an AiryScan detector. Ovary germline nuclei from stage 6-9 egg chambers were imaged separately with a ×63 oil lens in a *Z*-stack of 120 planes of 150 nm intervals.

### Yeast Two-Hybrid Analysis

Yeast two-hybrid screening was performed by Hybrigenics Services, S.A.S., Paris, France (http://www.hybrigenics-services.com). The coding sequence for full-length *Drosophila melanogaster* deadlock (NCBI reference NM_136247.3) was PCR-amplified and cloned into pB27 as a C-terminal fusion to LexA (LexA-del). The construct was sequence verified and used as a bait to screen a random-primed *Drosophila* ovary cDNA library constructed into pP6. pB27 and pP6 derive from the original pBTM116 (Vojtek and Hollenberg, 1995) and pGADGH (Bartel et al., 1993) plasmids, respectively. 105 million clones (10-fold the complexity of the library) were screened using a mating approach with YHGX13 (Y187 ade2-101::loxP-kanMX-loxP, mat α) and L40ΔGal4 (mata) yeast strains as previously described (Fromont-Racine et al., 1997). 185 His+ colonies were selected on a medium lacking tryptophan, leucine and histidine, and supplemented with 1 mM 3-aminotriazole to handle bait auto-activation. The prey fragments of the positive clones were amplified by PCR and sequenced at their 5’ and 3’ junctions. The resulting sequences were used to identify the corresponding interacting proteins in the GenBank database (NCBI).

### Imaging data analysis for localization studies

Imaging data analysis was done as described in (Andersen et al., 2017) with modifications. Acquired images were deconvolved using Huygens Professional Software (Scientific Volume Imaging, SVI). Chromatic aberrations in every sample were measured using TetraSpeck beads and corrected using Chromatic Aberration Correction (CAC) macro which relied on the Descriptor-based registration plugin (Preibisch et al., 2010). The deconvolved and beads-aligned images were segmented using the DAPI and WGA signals into nuclear and nuage regions. Channel 1(*cluster 20A* RNA FISH in far-red channel) and Channel 2 (*cluster 42AB* RNA FISH in the red channel) in case of null mutant experiments and Channel 1(*cluster 42AB* RNA FISH in the red channel) and Channel 2 (GFP-tagged protein in the green channel) in case of protein localization experiments. For segmentation, a DoG filter was applied to the channels before thresholding. The center pixels of the loci were determined by finding local maxima using a Min/Max filter. These centers were then used as seed points for splitting compound objects as well as to calculate the distance of each focus to the prior found nuclear border. Size and intensity thresholds were applied to exclude ambiguous objects. Intensities, size, number, and distances per identified foci, cell and channel was exported for analysis and plotting in R.

### Defining and Curating 1 kb Genomic Windows

The genome 1kb tiles were generated as described by (Andersen et al., 2017; Mohn et al., 2014). Tile annotation and curation can be reproduced using the scripts available through (Andersen et al., 2017).

### ChIP-Seq library preparation

Chromatin immunoprecipitation (ChIP) was performed as previously described (Lee et al., 2006) with minor modifications. Briefly, 100 ovary pairs were dissected into ice-cold PBS and cross-linked in 1.8 % para-formaldehyde/PBS for 10 minutes at room temperature with agitation. The cross-linking reaction was quenched by addition of glycine and ovaries were washed in PBS and homogenized in a glass douncer with a tight pestle. The homogenates were then lysed on ice for 20 minutes. DNA was sheared using a Covaris E220 Ultrasonicator, sonicating each sample for 20 minutes at 4°C with 200 cycles per burst, duty factor at 5.0 and peak incident power (PIP) set to 140 W. The sonicated lysates were cleared by centrifugation and then incubated overnight at 4°C with antibodies specific to the target epitope. 50 µL 1:1 mix of Protein A and Protein G Dynabeads was then added and allowed to bind antibody complexes by incubation for 2 hours at 4°C. Following six washing steps, DNA-protein complexes were eluted and de-cross-linked overnight at 65°C. RNA and protein was digested by RNase A and Proteinase K treatment, respectively, before purification using ChIP DNA Clean & Concentrator columns (Zymo Research). Barcoded libraries were prepared using the NEBNext Ultra II DNA Library Prep Kit for Illumina (NEB), which were sequenced on a HiSeq2500 (Illumina).

### ChIP-Seq Analysis

ChIPseq reads were trimmed to high quality bases 5-45 before mapping to the *Drosophila melanogaster* genome (dm6, r6.18) using Bowtie (release 1.2.2) with 3-mismatch tolerance. Reads were then computationally extended to the library median insert length of 353 nt. Normalization between samples was done based on the number of genome-unique mapping reads for each sample. Subsequent quantification of reads mapping to 1 kb tiles was done using bedtools, while relative quantification and plotting was done in R (see *Data visualization* below). Briefly, Rhino ChIP-seq tile signal was normalized to corresponding values for the relevant ChIP input libraries. A pseudo-count of 1 was then added to each tile value before calculation of log2 fold-enrichment values relative to input.

### RNA-Seq library preparation

For RNAseq libraries from samples depleted of rRNA by RNaseH-treatment (RH RNAseq), we modified the protocol published in (Morlan et al., 2012). TRIzol-isolated total RNA was purified by RNAeasy columns with on-column DNase I digest (Qiagen) in accordance with manufacturer’s instructions. Depletion of rRNA from the purified total RNA was done by using a mix of antisense oligonucleotides matching *Drosophila melanogaster* rRNAs (listed in Supplementary Table 5). The oligonucleotides were added to the 1 µg RNA in RNase H Buffer (20 mM Tris-HCl pH=8, 100 mM NaCl) and annealed with a temperature gradient from 95 °C to 45 °C. The RNA-DNA hybrids were specifically digested with Hybridase Thermostable RNase H (Epicentre) at 45 °C for 1 hour. Next, DNA was digested with TURBO DNase (Invitrogen) and RNA was purified using RNA Clean & Concentrator-5 (Zymo) according to the manufacturer’s instructions. Libraries were then cloned using the NEBNext Ultra II Directional RNA Library Prep Kit for Illumina (NEB), following the recommended kit protocol and sequenced on a HiSeq2500 (Illumina).

PolyA-enriched RNAseq libraries were prepared in an automated setup on a Biomek i7 robot. Briefly, 0.5 µg total RNA was denatured at 72 °C for 3 minutes, then put on ice. PolyA+ enrichment was done with the NEBNext Poly(A) mRNA Magnetic Isolation Module (NEB) according to the manufacturer’s recommendations. DNA was prepared from polyA+ RNA using NEB Ultra II Directional First and Second Strand Synthesis modules (NEB) according to the manufacturer’s recommendations, but incubating only 7 minutes at 94 °C. The cDNA was purified with 1.5× AmpureXP beads and subjected to NEB UltraII DNA library preparation Kit. PCR fragments of 200-800 bp were purified and sequenced on a NextSeq550 instrument (Illumina).

### RNA-Seq Analysis

RNA-Seq reads were trimmed to high quality bases 5-45 before mapping to the *Drosophila* genome (dm6, r6.18) using STAR (Dobin et al., 2013) or to *Drosophila melanogaster* mRNA and transposon and host mRNA sequences using Salmon (0.10.2) (Patro et al., 2017). For genomic mapping by STAR, normalization between samples was done based on the number of genome-unique mapping reads for each sample. Subsequent quantification of reads mapping to 1 kb tiles (performed for RH RNAseq only) was done using bedtools, while relative quantification and plotting was done in R (see *Data visualization* below). Briefly, RNA-seq tile signal was normalized to the estimated mappability scores for each 1 kb window. A pseudo-count of 1 was then added to each tile value before calculation of log2 fold-change values relative to control genotype samples. For analyses of transposon and host mRNA levels, only poly-A-enriched RNAseq libraries were used in order to avoid confounding the analyses with non-polyadenylated piRNA precursor transcripts. For each library, transcripts-per-million-transcripts (TPM) values were calculated using Salmon (0.10.2). TPMs were inter-sample-normalized using Sleuth (0.30.0) (Pimentel et al., 2017).

### Small RNA-Seq library preparation

Small RNA libraries were generated as described by (Jayaprakash et al., 2011) with modifications. Briefly, 18-29 nt long small RNAs were purified by preparative PAGE from 10 µg of total ovarian RNA. Following, the 3’ linker (containing four random nucleotides) was ligated overnight using T4 RNA Ligase 2, truncated K227Q (NEB) after which the products were recovered by a second PAGE purification. 5’ RNA linkers with four terminal random nucleotides were then ligated to the small RNAs using T4 RNA ligase (NEB) followed by an SPRI magnetic bead purification (in-house-produced beads similar to Agencourt RNAclean XP). The cloned small RNAs were then reverse transcribed and PCR amplified before sequencing on a HiSeq2500 (Illumina). Cloning oligo sequences are given in Supplementary Table 2.

### Small RNA-Seq Analysis

Single-end, 50-nucleotide small RNA sequencing reads were trimmed by removal of the 3’ linker sequence (AGATCGGAAGAGCACACGTCT), as well as the four random nucleotides at each end. Trimmed reads were mapped to the *Drosophila* genome (dm6, r6.18) using Bowtie (release 1.2.2) with 0-mismatch tolerance. Genome coverage was calculated and normalized to the number of uniquely mapping microRNA reads (in millions). Reads mapping to rRNA, tRNA, snRNA and snoRNA were excluded. Subsequent quantification of reads mapping to 1 kb tiles was done using bedtools, while relative quantification and plotting was done in R (see *Data visualization* below). Briefly, small RNA-seq tile signal was normalized to the estimated mappability scores for each 1 kb window. A pseudo-count of 1 was then added to each tile value before calculation of log2 fold-change values relative to control genotype samples.

### RIP-Seq library preparation

RIPseq protocol was adapted from (Zhang et al., 2012) with modifications. 80-100 ovaries from well fed flies (3-5 days old) were dissected in ice-cold PBS and homogenized by 20 pestle strokes in RIP lysis buffer (50mM HEPES pH 7.3, 150mM KCL, 3.2 mM MgCl, 0.5% TERGITOL solution (Sigma)) freshly supplemented with RNaseOUT (2.5µL/mL), 1mM Pefabloc, cOmplete Protease Inhibitor Cocktail (Roche), and 0.5 mM DTT. The lysate was sonicated using Bioruptor at medium strength (15 seconds ON, 60 seconds OFF), and cleared by centrifugation at 14,000 rpm for 15 minutes at 4°C. 5% of cleared lysate were set aside to serve as input samples, while the remainder was incubated at 4°C with 30 µL anti-FLAG M2 magnetic beads for 3 hours. The beads were washed 2 times with RIP lysis for 10 minutes, transferred into a fresh tube, and washed for 2 more times with all washes performed at 4°C. RNA was extracted from input lysate and beads using TRIzol reagent (Invitrogen) according to manufacturer’s instructions, further purified using the RNA Clean & Concentrator −5 kit (Zymo Research) with In-Column DNase I treatment, and eluted in 8 µL nuclease free water. Half of the eluted RNA was used for strand-specific RIP-Seq library preparation using the NEBNext Ultra II Directional RNA Library Prep Kit (New England Biolabs).

### RIP-qPCR

Enrichment of cluster transcripts after RIP was assayed by RT-qPCR. cDNA was synthesized from 2 µL of purified and DNase treated RIP-RNA using the Maxima First Strand cDNA Synthesis Kit (Thermo Fisher Scientific) according to manufacturer’s protocol. cDNAs were amplified using primers listed in Supplementary Table 2. qPCR reactions were performed using GoTaq qPCR system (Promega).

### RIP-Seq analyses

RIP-Seq reads were trimmed to high quality bases 5-45 before mapping to the *Drosophila* genome (dm6, r6.18) using STAR (Dobin et al., 2013). Normalization between samples was done based on the number of high-quality reads sequenced for each sample. Subsequent quantification of reads mapping to 1 kb tiles was done using bedtools, while relative quantification and plotting was done in R (see *Data visualization* below). Briefly, RIP-seq tile signal was normalized to corresponding values for the relevant RIP input libraries. A pseudo-count of 1 was then added to each tile value before calculation of log2 fold-enrichment values relative to control RIP libraries from immunoprecipitation of NLS-GFP-V5-3xFLAG fusion proteins expressed in ovary nurse cells.

### PRO-Seq library preparation

PRO-seq was performed as described previously (Mahat et al., 2016), with modifications. Briefly, for each genotype, 50 µL ovaries were dissected into ice-cold PBS. Ovaries were then fragmented by douncing using a tight glass pestle and the homogenates were permeabilized and crude nuclei fractions were prepared by washing in permeabilization buffer. The permeabilized ovary samples were then stored at −80°C in *storage buffer* until further processing. Nuclear run-on reactions were performed with biotin-11-CTP and biotin-11-UTP and unlabeled ATP and GTP. Following RNA purification from the run-on reactions, RNA samples were fragmented for 10 minutes on ice in 0.2 N NaOH and purified on Bio-Spin P30 columns (Bio-Rad). Biotin-labelled RNA was then purified using MyOne Streptavidin C1 Dynabeads (Thermo Fisher Scientific) and decapped using Tobacco Acid Pyrophosphatase (TAP) enzyme (Epicentre, can be replaced by RppH from NEB) and purified by phenol/chloroform extraction. 3’ linkers and then 5’ linkers (same as for small RNA cloning) were then ligated subsequently and biotinylated ligation products were purified after each ligation using MyOne Streptavidin C1 Dynabeads. Reverse transcription was performed using SuperScript II enzyme (Thermo Fisher Scientific) and the cloned libraries were PCR amplified using KAPA Hotstart Realtime master mix (KAPA Biosystems). The amplified libraries were then sequenced on a HiSeq2500 (Illumina) in paired-end 75 bp mode.

### PRO-Seq analysis

Second mate-pair reads, representing RNA 3’ ends, were mapped to the *Drosophila* genome (dm6, r6.18) using STAR (Dobin et al., 2013). Normalization between samples was done based on the number of genome-unique mapping reads for each sample. Subsequent quantification of reads mapping to 1 kb tiles was done using bedtools, while relative quantification and plotting was done in R (see *Data visualization* below). Briefly, PRO-Seq tile signal was normalized to the estimated mappability scores for each 1 kb window. A pseudo-count of 1 was then added to each tile value before calculation of log2 fold-change values relative to control genotype samples.

### NXF family protein sequence alignments

Sets of sequences orthologous to *D. melanogaster* Sbr (Nxf1) and Nxf3 were collected using reciprocal protein blast searches. A multiple sequence alignment of the selected sequences was generated using mafft (v7.407) and visualized using Jalview (v2.10.2b2) with the clustal color-scheme. All sequence accession numbers are given in Supplementary Table 6.

### Data visualization

Data visualization and statistical analyses were done using R (RCoreTeam, 2018) in conjunction with the following software packages: ggplot2 (Wickham, 2016), tidyverse (https://www.tidyverse.org/) and scales (Wickham, 2017). The UCSC genome browser (Kent et al., 2002; Raney et al., 2013) was used to explore sequencing data and to prepare genome browser panels shown in the manuscript.

## DATA AVAILABILITY

All sequencing data produced for this publication has been deposited to the NCBI GEO archive under the accession number GSE126578.

The mass spectrometry proteomics data have been deposited to the ProteomeXchange Consortium via the PRIDE (Vizcaino et al., 2016) partner repository with the dataset identifier PXD012717.

---

**Figure S1.**
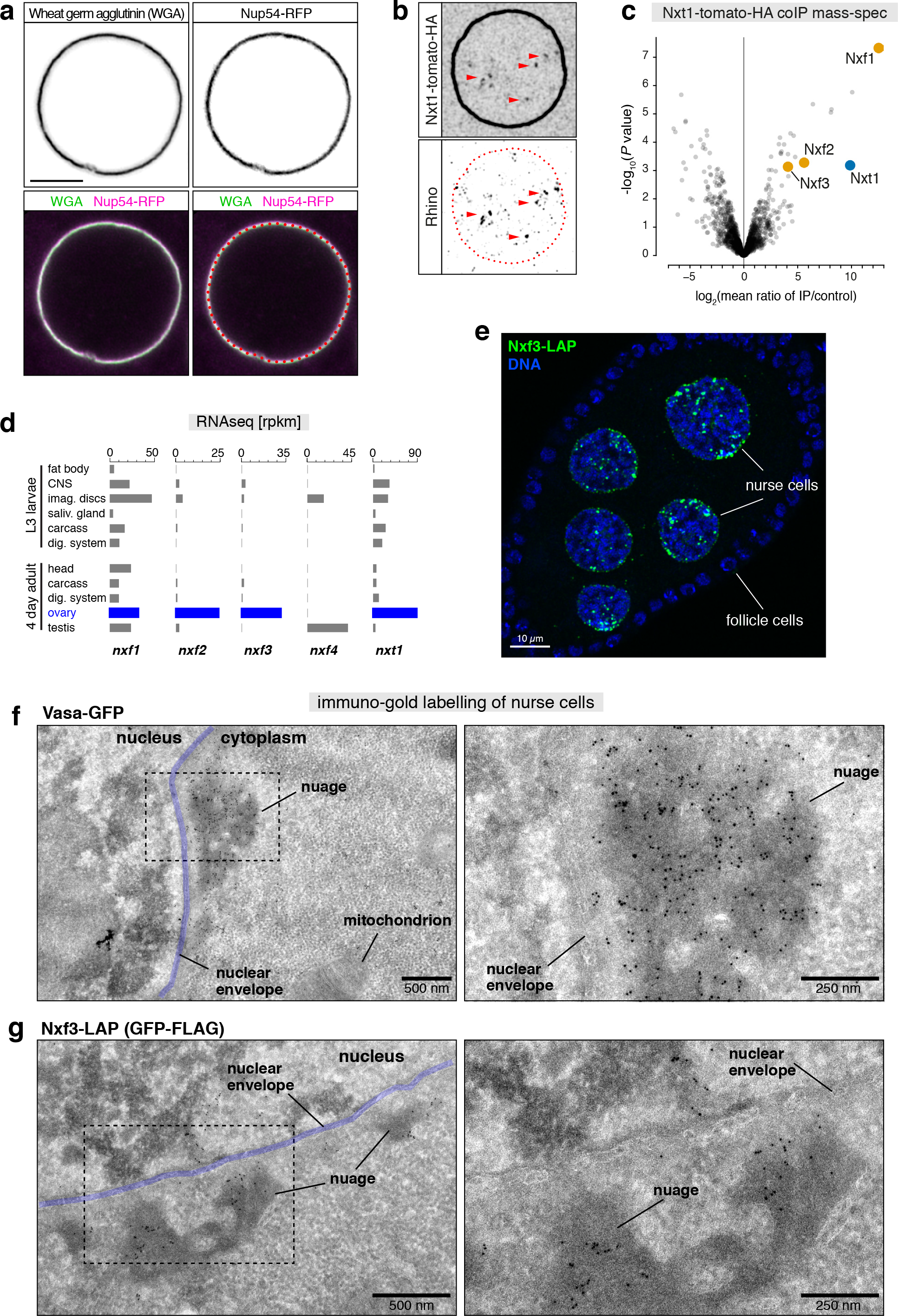
(related to Figure 1). **a**, Confocal images of a nurse cell nucleus (~stage 8 egg chamber; scale bar: 5 μm) expressing Nup54-RFP under its endogenous promoter region, stained for nuclear pore complexes with wheat germ agglutinin (WGA). The WGA signal overlaps precisely with that of Nup58-RFP. Throughout the paper, we used WGA to visualize the nuclear envelope (red dotted line corresponds to center of WGA signal; lower right panel). **b**, Localization of Nxt1-tdTomato-HA alongside Rhino in nurse cells (red dotted line: nuclear envelope; red arrowheads: Rhino foci; scale bar: 5 μm). **c**, Volcano plot showing enrichment values and corresponding significance levels for proteins co-purifying with Nxt1-tdTomato-HA from ovaries versus control (Wild type ovaries; n = 3). **d**, mRNA levels as determined by RNAseq for indicated genes from indicated larval or adult tissues (data from FlyBase). **e**, Confocal image of an entire egg chamber expressing Nxf3-LAP from its endogenous locus (nurse cell and follicle cell nuclei are indicated; scale bar: 10 μm). **f, g**, Transmission electron microscopy images of nurse cells at different magnifications. The nuclear envelope is pseudo-colored (blue), also indicated are mitochondria, nuage (electron-dense perinuclear structures), nucleus and cytoplasm. Sections were stained with a polyclonal anti-GFP antibody and a secondary anti-rabbit antibody coupled to 6nm gold particles (f, ovaries expressing Vasa-GFP; g, ovaries expressing Nxf3-GFP).

**Figure S2.**
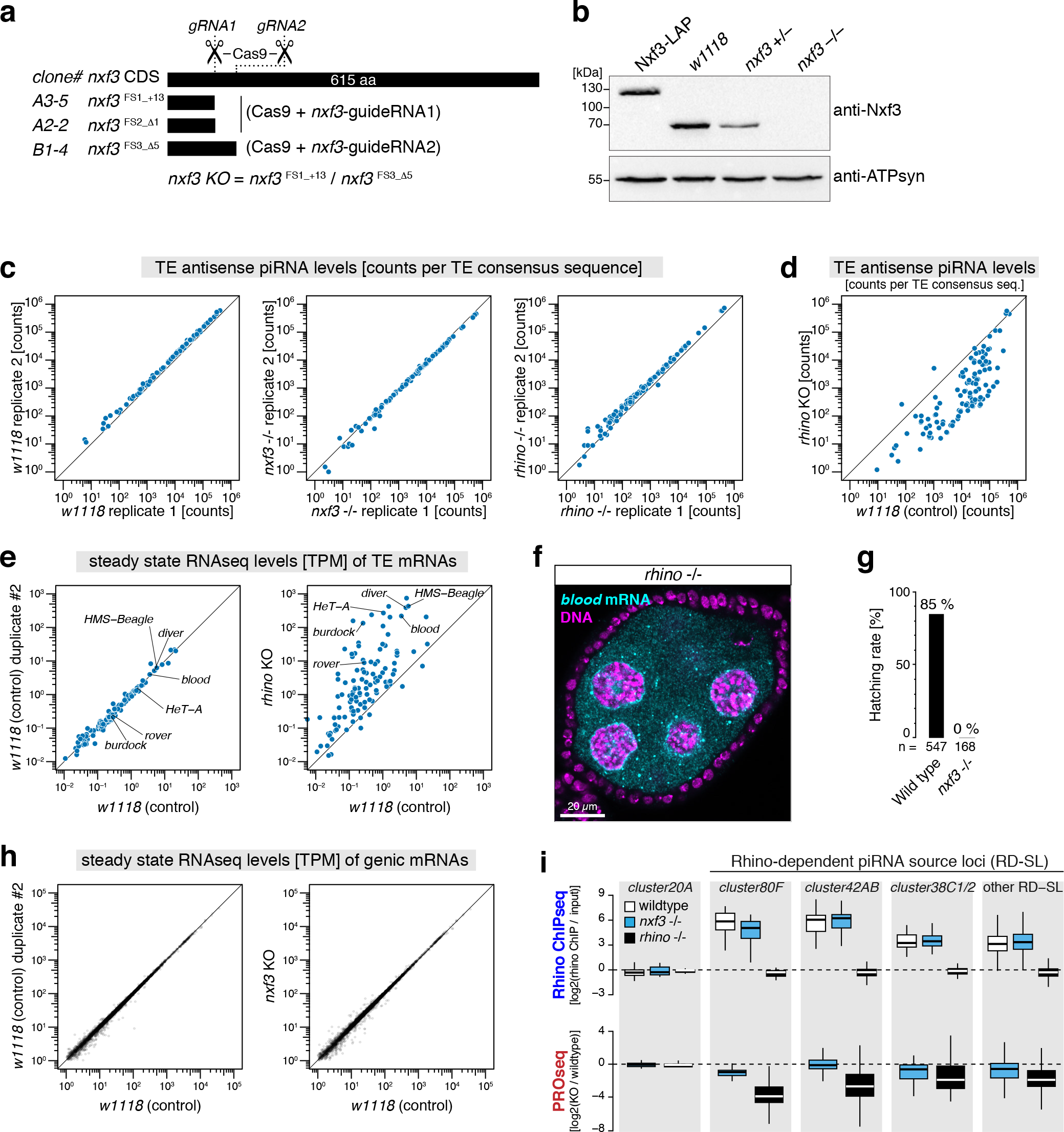
(related to Figure 2). **a**, Cartoon depicting the frameshift mutations in nxf3 generated by CRISPR/Cas9 using two independent single guide RNAs. The allelic null allele combination used through-out this paper is indicated at the bottom. **b**, Western blot analysis using a monoclonal antibody raised against Nxf3 verifying the endogenous LAP-tagging at the nxf3 locus (lane 1), and the nxf3 mutant frameshift alleles (lane 4). A staining against ATP-synthetase serves as loading control. **c**, Scatter plots displaying levels of antisense piRNAs mapping to consensus transposon sequences from ovary samples of the two replicates per genotype. **d**, Scatter plot displaying levels of antisense piRNAs mapping to consensus transposon sequences from control ovary samples versus rhino mutant samples. **e**, Scatter plots displaying levels of sense RNA reads mapping to consensus transposon sequences from ovary samples of indicated genotype. **f**, Confocal image of rhino mutant egg chamber showing the blood transposon mRNA (FISH: cyan; DAPI: magenta). Image taken under identical settings as those shown in Figure 2f. **g**, Bar plot indicating embryo hatching rate of eggs laid by w[1118] control females and nxf3 mutant females. **h**, Scatter plots displaying steady state mRNA levels (transcripts per million transcripts, TPM) from ovary samples of indicated genotype. **i**, Boxplot displaying log2 fold changes in Rhino occupancy (top panel; ChIP-seq) or transcriptional output (bottom panel; PRO-seq) at indicated genomic loci in ovaries of indicated genotypes. Contrasted is the Rhino-independent piRNA cluster20A against Rhino-dependent piRNA clusters (42AB, 80F, 38C1+2, others). Individual data points underlying the boxes are genomic 1kb tiles, only reads mapping uniquely to each locus were considered. Boxplot definition as in Figure 2b.

**Figure S3.**
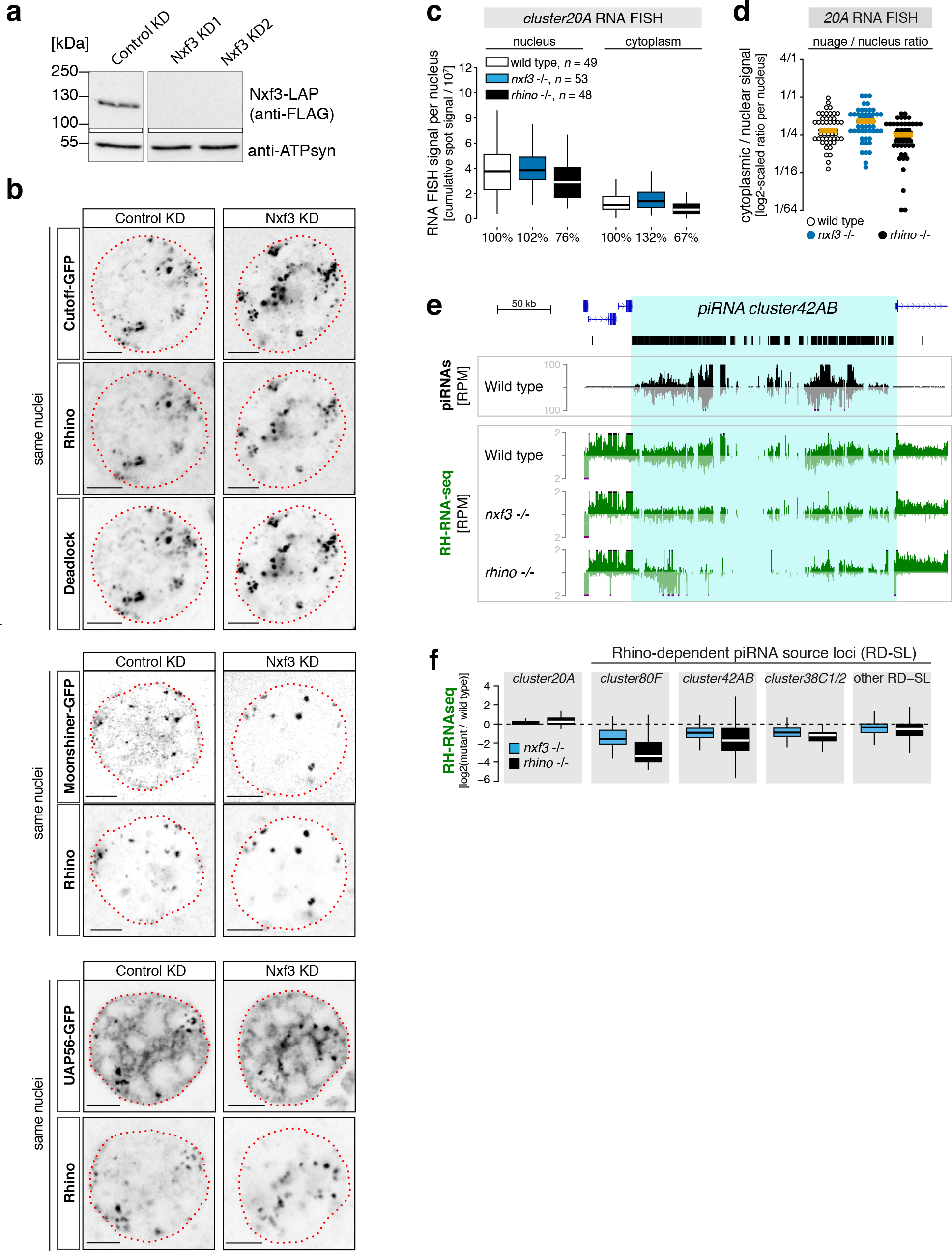
(related to Figure 2). **a**, Western blot verifying the efficiency of two independent sh-RNA lines against Nxf3. sh-RNA lines were expressed with MTD-Gal4 in ovaries expressing Nxf3-LAP. A staining against ATP-synthetase serves as loading control. **b**, Confocal images confirming that loss of Nxf3 (depleted via sh-RNA expression) does not impact enrichment of Cutoff, Rhino, Deadlock, Moonshiner, or UAP56 at nuclear Rhino foci (piRNA clusters). **c**, Quantification of cluster20A RNA FISH signal in nurse cells with indicated genotype (boxplot definition as in Figure 2b; number of analyzed germline nuclei indicated). **d**, Ratio of total cluster20A RNA FISH signal in nuage and nucleus quantified per nurse cell nucleus (genotypes indicated; orange bars: median values). **e**, UCSC genome browser panel showing ovarian steady state RNA levels (reads per million reads; total RNA-seq after depletion of rRNA) at cluster42AB in indicated genotypes. **f**, Boxplot displaying log2 fold changes in steady state RNA levels (RNA-seq) at indicated genomic loci in ovaries of indicated genotypes (respective to control samples). Contrasted is the Rhino-inde-pendent piRNA cluster20A against Rhino-dependent piRNA clusters (42AB, 80F, 38C1+2, others). Individual data points underlying the boxes are genomic 1kb tiles, only reads mapping uniquely to each locus were considered. Boxplot definition as in Figure 2b.

**Figure S4.**
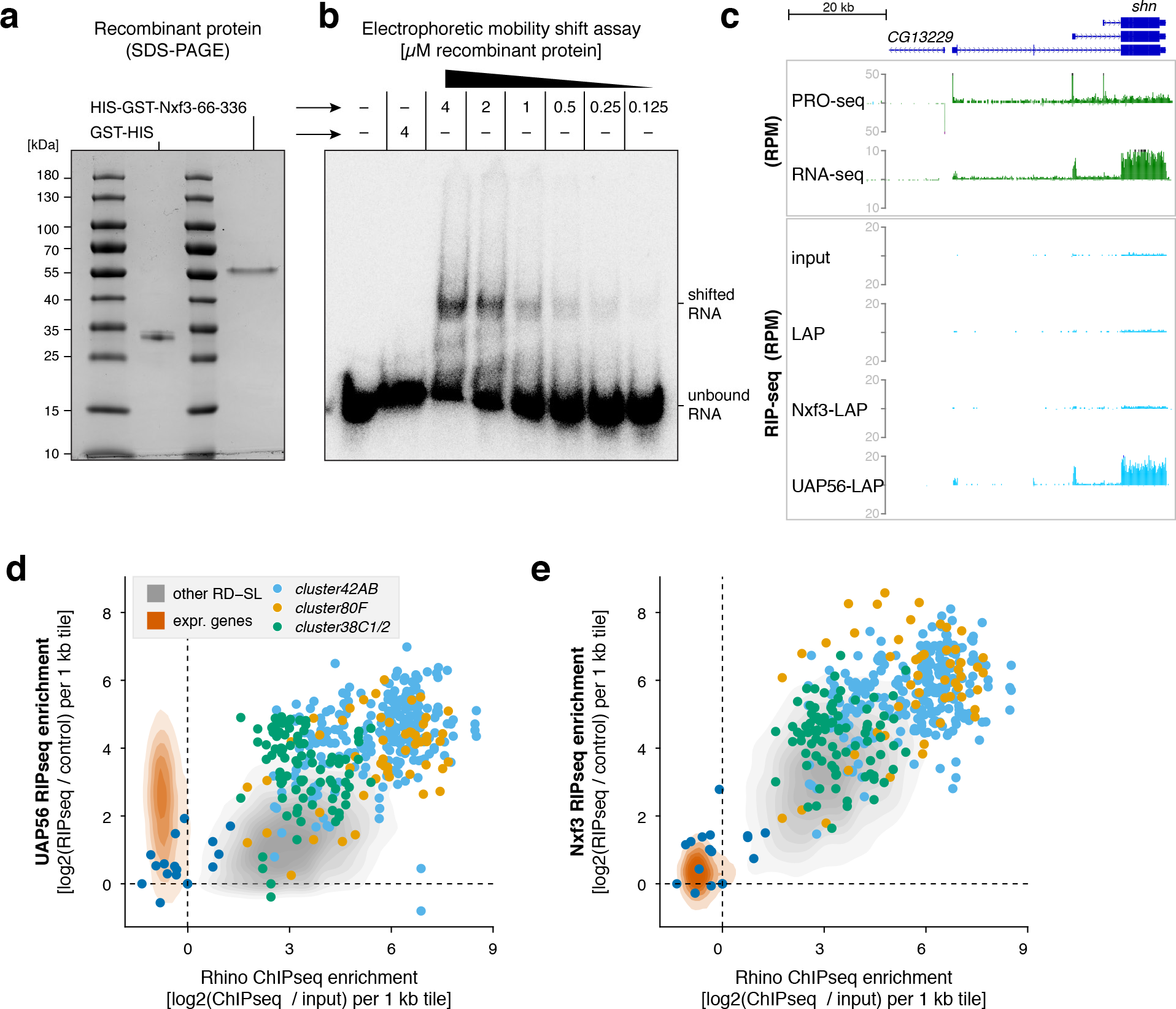
(related to Figure 3). **a**, Coomassie stained PAGE gel showing purity of recombinant GST-tagged Nxf3 RNA binding domain (residues 66-336). **b**, Native PAGE gel displaying gel shift of a radioactively labelled single-stranded RNA oligo upon addition of different amounts of the recombinant Nxf3 RNA binding domain. **c**, UCSC genome browser panel showing transcriptional output (PRO-seq), steady state RNA levels (RNA-seq) and RIP-seq signal in ovaries of indicated genotype at the schnurri locus. **d, e**, Scatter plots showing RIP-seq enrichment for UAP56 (d) or Nxf3 (e) versus Rhino occupancy. Contrasted are genomic 1kb tiles mapping to Rhino-dependent piRNA source loci (grey; major clusters shown as colored dots) versus genomic tiles overlapping expressed protein-coding genes (orange). Genomic 1kb tiles belonging to the Rhino-independent piRNA cluster20A are shown in dark blue.

**Figure S5.**
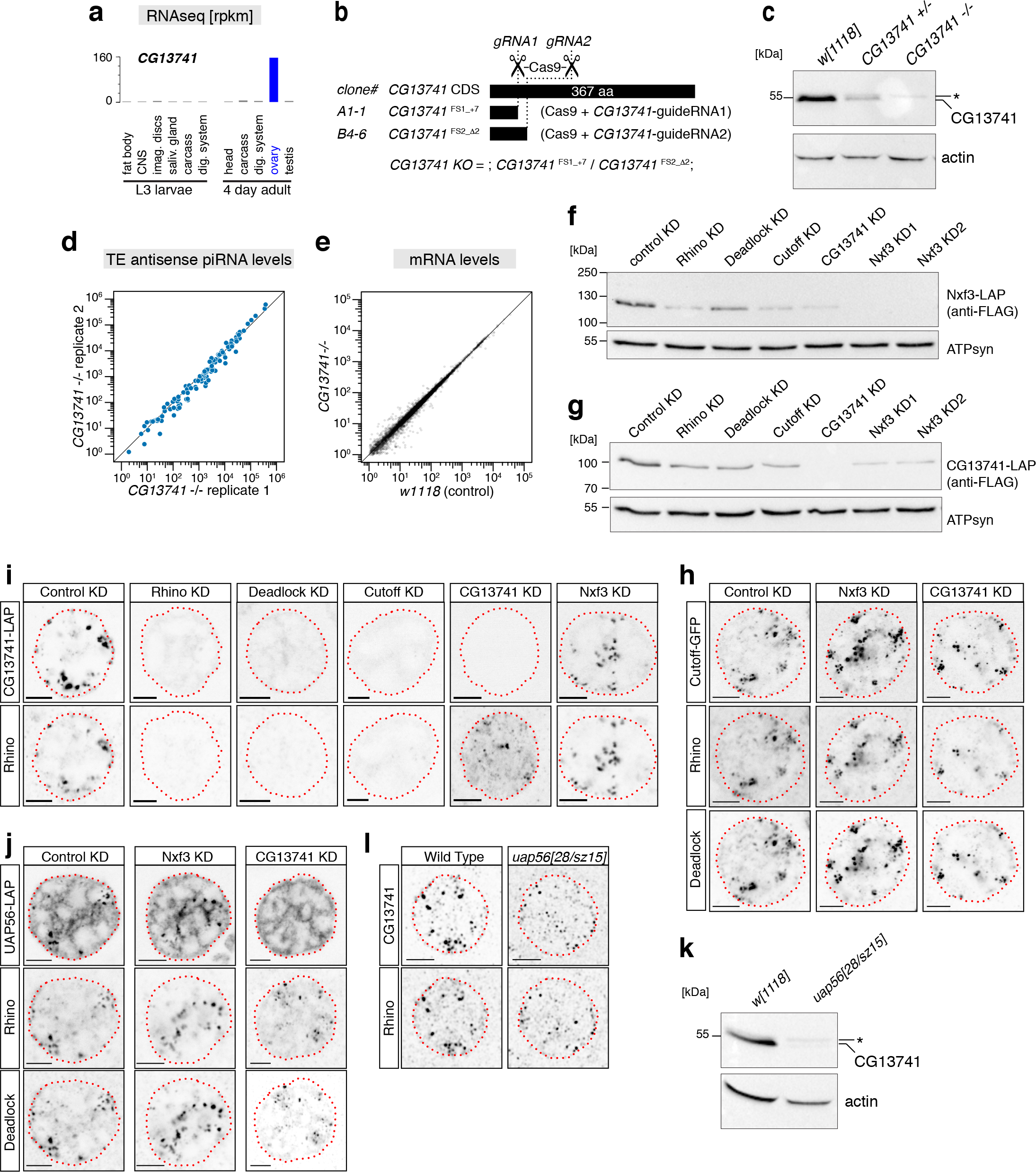
(related to Figure 4 and 5). **a**, Shown are mRNA levels as determined by RNAseq for CG13741 from indicated larval or adult tissues (data from Flybase). **b**, Cartoon depicting the frameshift mutations in CG13741 generated by CRISPR/Cas9 using two independent single guide RNAs. The allelic null allele combination used throughout this paper is indicated at the bottom. **c**, Western blot analysis using a monoclonal antibody raised against CG13741 verifying the CG13741 mutant frameshift alleles (the asterisk marks an unspecific band). A staining against actin served as loading control. **d**, Scatter plot displaying levels of antisense piRNAs mapping to consensus transposon sequences from the two replicate CG13741 mutant samples. **e**, Scatter plot displaying steady state mRNA levels (transcripts per million sequenced transcripts) from ovarian samples of indicated genotype. **f, g**, Western blots showing levels of endogenously tagged Nxf3-LAP (f) or CG13741-LAP (g) in lysates from ovaries of indicated genotype (KD: germline specific sh-RNA mediated knockdown). A staining against ATP-synthetase served as loading control. **h**, Confocal images of nurse cell nuclei (each column is the same nucleus; genotypes indicated at top) displaying localization of Cutoff-GFP, Rhino, and Deadlock (scale bars: 5 μm; red dotted line: nuclear envelope). **i**, Confocal images of nurse cell nuclei (each column is the same nucleus) displaying localization of CG13741-LAP and Rhino (scale bars: 5μm; red dotted line: nuclear envelope). **j**, Confocal images of nurse cell nuclei (each column is the same nucleus; genotypes indicated at top) displaying localization of UAP56-LAP, Rhino, and Deadlock (scale bars: 5 μm; red dotted line: nuclear envelope). **k**, Western blot analysis showing CG13741 levels in control and uap56 mutant ovary lysates (the asterisk marks an unspecific band). A staining against actin served as loading control. **l**, Confocal images of a uap56[28/sz15] mutant nurse cell indicating sub-cellular localization of CG13741-LAP and Rhino (scale bar: 5 μm; red dotted line: nuclear envelope).

**Figure S6.**
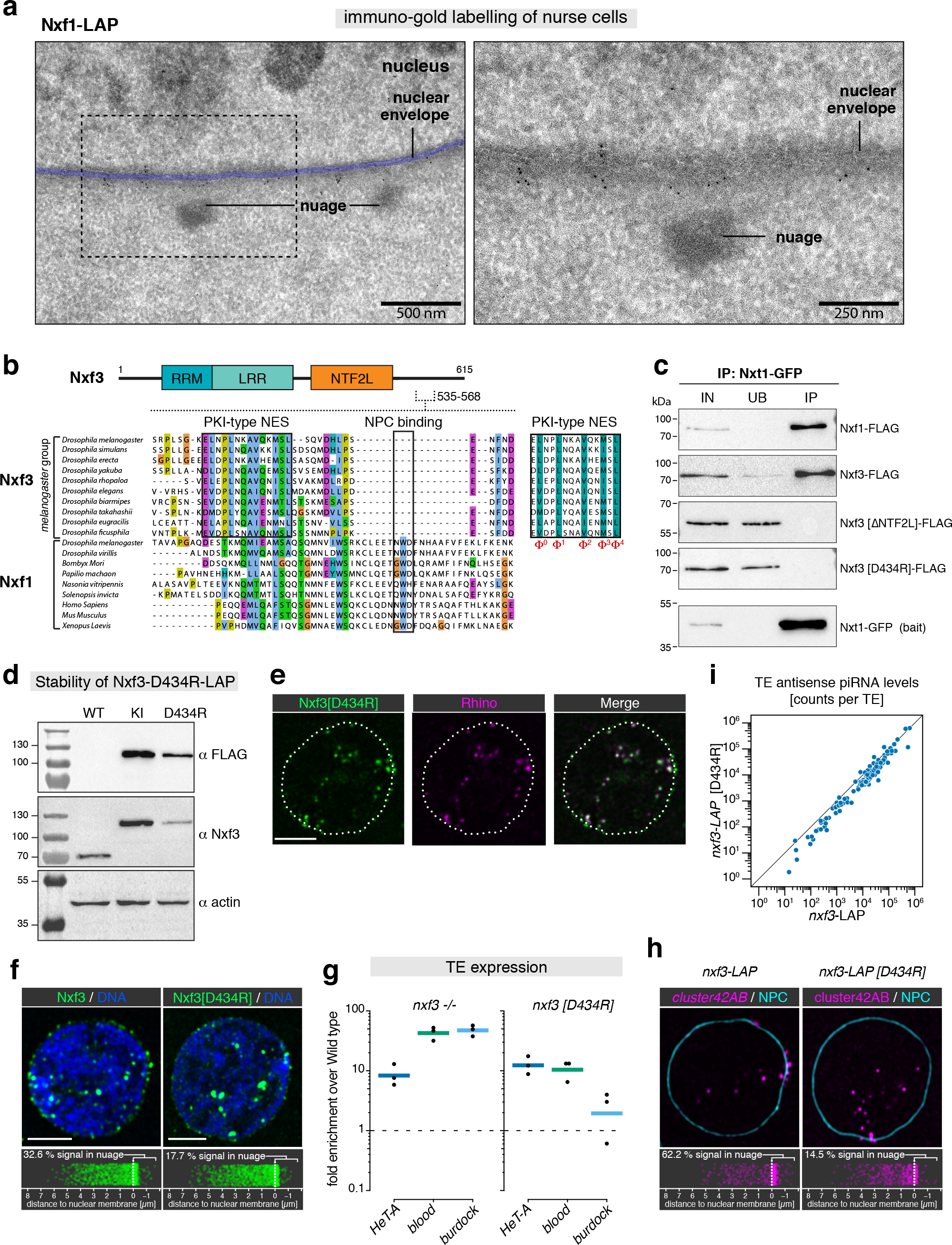
(related to Figure 6). **a**, Transmission electron microscopy images of nurse cells expressing Nxf1-LAP at different magnifications. The nuclear envelope is pseudo-colored (blue), also indicated are nuage (electron-dense perinuclear structures), and nucleus. Sections were stained with a polyclonal anti-GFP antibody and a secondary anti-rabbit antibody coupled to 6 nm gold particles. **b**, Protein sequence alignment displaying the nucleoporin interacting motif in Nxf1 proteins, as well as the position of the PKI-type NES motif in Nxf3 proteins. The NES core features (hydrophobic residues required for Crm1 interaction) are shown to the right. **c**, Western blot analysis of Nxt1-GFP co-immuno-precipitation experiments where indicated FLAG-tagged Nxf1 or Nxf3 constructs were co-transfected in S2 cells. **d**, Western blots indicating levels of endogenous Nxf3 (lane 1) and endogenously tagged Nxf3-LAP (lane 2: wildtype Nxf3-LAP; lane 3: Nxt1 interacting mutant Nxf3-LAP). **e**, Confocal images of a nurse cell nucleus (scale bar: 5 μm; white dotted line: nuclear envelope) showing co-localization of Nxf3-LAP [D434R] (green) and Rhino (magenta). **f**, Confocal images of representative nurse cell nuclei (scale bar: 5 μm) showing localization of indicted Nxf3-LAP proteins (DAPI: blue). Jitter plots below indicate the distance of Nxf3-LAP protein foci to the nuclear envelope (n = 1500 foci analyzed per genotype). **g**, Shown are changes in steady state RNA levels of indicated transposons (as determined via quantitative RT-PCR) in ovaries of indicated genotype versus Wild type control (n = 3; bar marks mean). **h**, As in (f), but instead of Nxf3 signal, cluster42AB FISH signal is shown (same nucleus as in (f) is shown).

**Figure S7.**
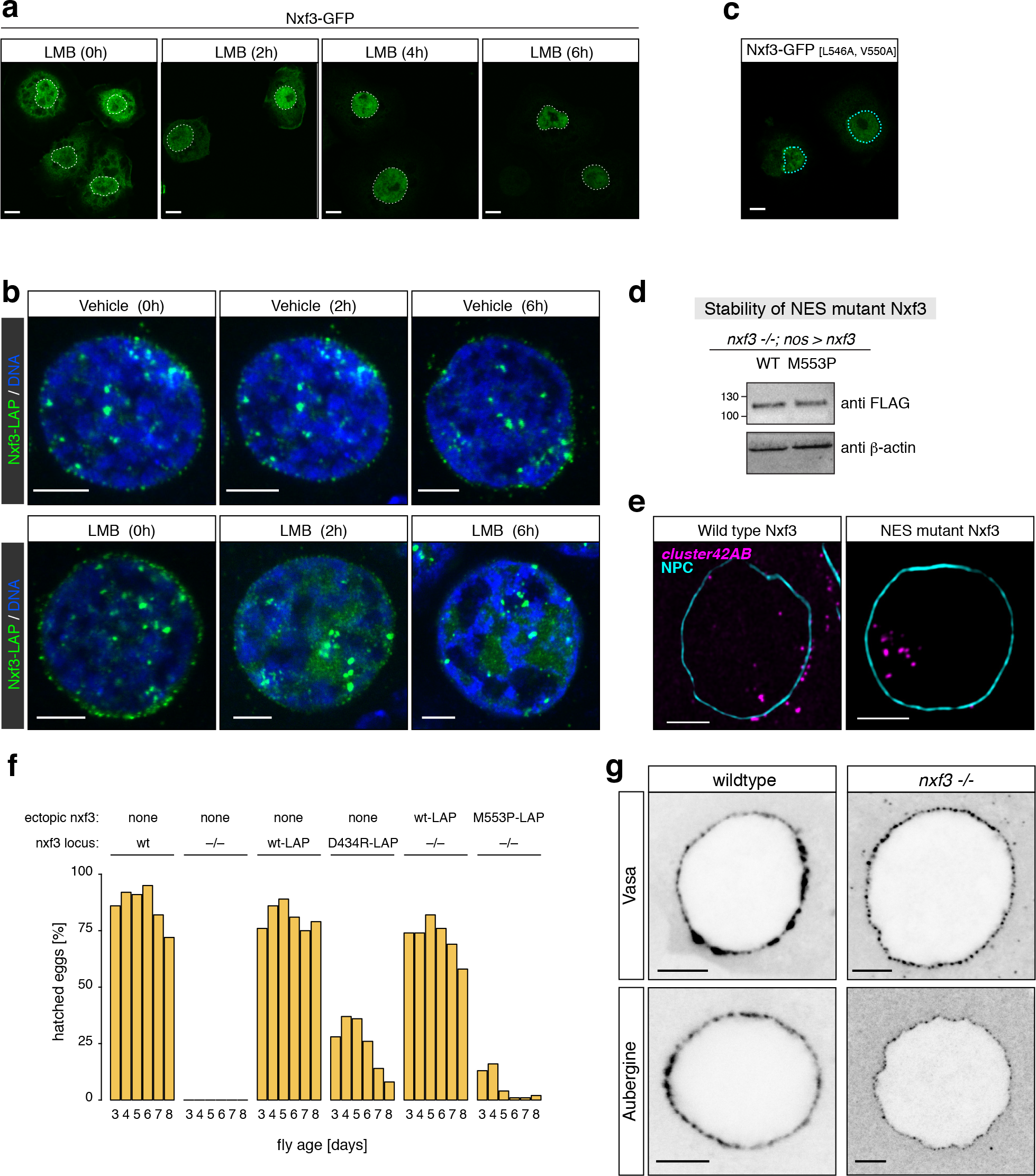
(related to Figure 6). **a**, Confocal images showing sub-cellular localization of Nxf3-GFP in S2 cells treated for indicated times with the Crm1 inhibitor LMB (scale bar: 5μm; dotted line marks nuclear envelope). **b**, Confocal images showing sub-cellular localization of Nxf3-LAP in ovaries treated for indicated times with the Crm1 inhibitor LMB (bottom row) or carrier (7-methanol; top row; scale bar: 5 μm). Note the loss of Nxf-LAP signal in nuage upon LMB treatment. **c**, Confocal image showing sub-cellular localization of Nxf3-GFP [L546A, V550A] in S2 cells (scale bar: 5 μm). **d**, Western blot analysis showing levels of nanos promoter driven Nxf3-LAP (Wild type versus NES mutant) in ovaries mutant for endogenous nxf3. **e**, Confocal images (scale bar: 5 μm) of nurse cells expressing Wild type Nxf3-LAP (left) or Nxf3-LAP [M553P] (right) stained for cluster42AB transcripts (magenta) alongside nuclear pores (WGA: cyan). Example image for the jitter plot shown in Figure 6g. **f**, Bar plot summarizing hatching rate (in percent) of eggs laid by females of indicated age and genotype mated to wildtype males. **g**, confocal images of nurse cells (genotype indicated on top) expressing Vasa-GFP or Aubergine-GFP. Nuage granules are smaller, but clearly present in ovaries mutant for nxf3.

## REFERENCES

Aibara, S., Katahira, J., Valkov, E., and Stewart, M. (2015). The principal mRNA nuclear export factor NXF1:NXT1 forms a symmetric binding platform that facilitates export of retroviral CTE-RNA. Nucleic acids research 43, 1883–1893.

Andersen, P.R., Tirian, L., Vunjak, M., and Brennecke, J. (2017). A heterochromatin-dependent transcription machinery drives piRNA expression. Nature 549, 54–59.

Bohnsack, M.T., Czaplinski, K., and Gorlich, D. (2004). Exportin 5 is a RanGTP-dependent dsRNA-binding protein that mediates nuclear export of pre-miRNAs. Rna 10, 185–191.

Braun, I.C., Herold, A., Rode, M., and Izaurralde, E. (2002). Nuclear export of mRNA by TAP/NXF1 requires two nucleoporin-binding sites but not p15. Molecular and cellular biology 22, 5405–5418.

Brennecke, J., Aravin, A.A., Stark, A., Dus, M., Kellis, M., Sachidanandam, R., and Hannon, G.J. (2007). Discrete Small RNA-Generating Loci as Master Regulators of Transposon Activity in Drosophila. Cell 128, 1089–1103.

Brown, J.B., Boley, N., Eisman, R., May, G.E., Stoiber, M.H., Duff, M.O., Booth, B.W., Wen, J., Park, S., Suzuki, A.M., et al. (2014). Diversity and dynamics of the Drosophila transcriptome. Nature 512, 393–399.

Chen, Y.A., Stuwe, E., Luo, Y., Ninova, M., Le Thomas, A., Rozhavskaya, E., Li, S., Vempati, S., Laver, J.D., Patel, D.J., et al. (2016). Cutoff Suppresses RNA Polymerase II Termination to Ensure Expression of piRNA Precursors. Molecular cell 63, 97–109.

Cullen, B.R. (2000). Nuclear RNA export pathways. Molecular and cellular biology 20, 4181–4187.

Czech, B., Munafo, M., Ciabrelli, F., Eastwood, E.L., Fabry, M.H., Kneuss, E., and Hannon, G.J. (2018). piRNA-Guided Genome Defense: From Biogenesis to Silencing. Annual review of genetics 52, 131–157.

Czech, B., Preall, J.B., McGinn, J., and Hannon, G.J. (2013). A Transcriptome-wide RNAi Screen in the Drosophila Ovary Reveals Factors of the Germline piRNA Pathway. Molecular cell 50, 749–761.

Dennis, C., Brasset, E., Sarkar, A., and Vaury, C. (2016). Export of piRNA precursors by EJC triggers assembly of cytoplasmic Yb-body in Drosophila. Nature communications 7, 13739.

Dobin, A., Davis, C.A., Schlesinger, F., Drenkow, J., Zaleski, C., Jha, S., Batut, P., Chaisson, M., and Gingeras, T.R. (2013). STAR: ultrafast universal RNA-seq aligner. Bioinformatics 29, 15–21.

Doblmann, J., Dusberger, F., Imre, R., Hudecz, O., Stanek, F., Mechtler, K., and Durnberger, G. (2018). apQuant: Accurate Label-Free Quantification by Quality Filtering. J Proteome Res.

Dorfer, V., Pichler, P., Stranzl, T., Stadlmann, J., Taus, T., Winkler, S., and Mechtler, K. (2014). MS Amanda, a universal identification algorithm optimized for high accuracy tandem mass spectra. J Proteome Res 13, 3679–3684.

Durdevic, Z., Pillai, R.S., and Ephrussi, A. (2018). Transposon silencing in the Drosophila female germline is essential for genome stability in progeny embryos. Life Sci Alliance 1, e201800179.

Ejsmont, R.K., Bogdanzaliewa, M., Lipinski, K.A., and Tomancak, P. (2011). Production of fosmid genomic libraries optimized for liquid culture recombineering and cross-species transgenesis. Methods in molecular biology 772, 423–443.

Fribourg, S., Braun, I.C., Izaurralde, E., and Conti, E. (2001). Structural basis for the recognition of a nucleoporin FG repeat by the NTF2-like domain of the TAP/p15 mRNA nuclear export factor. Molecular cell 8, 645–656.

Gokcezade, J., Sienski, G., and Duchek, P. (2014). Efficient CRISPR/Cas9 Plasmids for Rapid and Versatile Genome Editing in Drosophila. G3.

Goriaux, C., Desset, S., Renaud, Y., Vaury, C., and Brasset, E. (2014). Transcriptional properties and splicing of the flamenco piRNA cluster. EMBO reports 15, 411–418.

Gratz, S.J., Cummings, A.M., Nguyen, J.N., Hamm, D.C., Donohue, L.K., Harrison, M.M., Wildonger, J., and O’Connor-Giles, K.M. (2013). Genome engineering of Drosophila with the CRISPR RNA-guided Cas9 nuclease. Genetics 194, 1029–1035.

Gruter, P., Tabernero, C., von Kobbe, C., Schmitt, C., Saavedra, C., Bachi, A., Wilm, M., Felber, B.K., and Izaurralde, E. (1998). TAP, the human homolog of Mex67p, mediates CTE-dependent RNA export from the nucleus. Molecular cell 1, 649–659.

Guttler, T., Madl, T., Neumann, P., Deichsel, D., Corsini, L., Monecke, T., Ficner, R., Sattler, M., and Gorlich, D. (2010). NES consensus redefined by structures of PKI-type and Rev-type nuclear export signals bound to CRM1. Nature structural & molecular biology 17, 1367–1376.

Han, B.W., Wang, W., Li, C., Weng, Z., and Zamore, P.D. (2015). Noncoding RNA. piRNA-guided transposon cleavage initiates Zucchini-dependent, phased piRNA production. Science 348, 817–821.

Heath, C.G., Viphakone, N., and Wilson, S.A. (2016). The role of TREX in gene expression and disease. The Biochemical journal 473, 2911–2935.

Herold, A., Klymenko, T., and Izaurralde, E. (2001). NXF1/p15 heterodimers are essential for mRNA nuclear export in Drosophila. Rna 7, 1768–1780.

Herold, A., Suyama, M., Rodrigues, J.P., Braun, I.C., Kutay, U., Carmo-Fonseca, M., Bork, P., and Izaurralde, E. (2000). TAP (NXF1) belongs to a multigene family of putative RNA export factors with a conserved modular architecture. Molecular and cellular biology 20, 8996–9008.

Herold, A., Teixeira, L., and Izaurralde, E. (2003). Genome-wide analysis of nuclear mRNA export pathways in Drosophila. The EMBO journal 22, 2472–2483.

Homolka, D., Pandey, R.R., Goriaux, C., Brasset, E., Vaury, C., Sachidanandam, R., Fauvarque, M.O., and Pillai, R.S. (2015). PIWI Slicing and RNA Elements in Precursors Instruct Directional Primary piRNA Biogenesis. Cell reports 12, 418–428.

Hur, J.K., Luo, Y., Moon, S., Ninova, M., Marinov, G.K., Chung, Y.D., and Aravin, A.A. (2016). Splicing-independent loading of TREX on nascent RNA is required for efficient expression of dual-strand piRNA clusters in Drosophila. Genes & development 30, 840–855.

Jayaprakash, A.D., Jabado, O., Brown, B.D., and Sachidanandam, R. (2011). Identification and remediation of biases in the activity of RNA ligases in small-RNA deep sequencing. Nucleic acids research 39, e141.

Katahira, J., Dimitrova, L., Imai, Y., and Hurt, E. (2015). NTF2-like domain of Tap plays a critical role in cargo mRNA recognition and export. Nucleic acids research 43, 1894–1904.

Katahira, J., Straesser, K., Saiwaki, T., Yoneda, Y., and Hurt, E. (2002). Complex formation between Tap and p15 affects binding to FG-repeat nucleoporins and nucleocytoplasmic shuttling. The Journal of biological chemistry 277, 9242–9246.

Kent, W.J., Sugnet, C.W., Furey, T.S., Roskin, K.M., Pringle, T.H., Zahler, A.M., and Haussler, D. (2002). The human genome browser at UCSC. Genome research 12, 996–1006.

Kerkow, D.E., Carmel, A.B., Menichelli, E., Ambrus, G., Hills, R.D., Jr., Gerace, L., and Williamson, J.R. (2012). The structure of the NXF2/NXT1 heterodimeric complex reveals the combined specificity and versatility of the NTF2-like fold. J Mol Biol 415, 649–665.

Klattenhoff, C., Xi, H., Li, C., Lee, S., Xu, J., Khurana, J.S., Zhang, F., Schultz, N., Koppetsch, B.S., Nowosielska, A., et al. (2009). The Drosophila HP1 homolog Rhino is required for transposon silencing and piRNA production by dual-strand clusters. Cell 138, 1137–1149.

Kohler, A., and Hurt, E. (2007). Exporting RNA from the nucleus to the cytoplasm. Nature reviews Molecular cell biology 8, 761–773.

Kudo, N., Matsumori, N., Taoka, H., Fujiwara, D., Schreiner, E.P., Wolff, B., Yoshida, M., and Horinouchi, S. (1999). Leptomycin B inactivates CRM1/exportin 1 by covalent modification at a cysteine residue in the central conserved region. Proceedings of the National Academy of Sciences of the United States of America 96, 9112–9117.

Kwak, H., Fuda, N.J., Core, L.J., and Lis, J.T. (2013). Precise maps of RNA polymerase reveal how promoters direct initiation and pausing. Science 339, 950–953.

Lee, T.I., Johnstone, S.E., and Young, R.A. (2006). Chromatin immunoprecipitation and microarray-based analysis of protein location. Nature protocols 1, 729–748.

Levesque, L., Guzik, B., Guan, T., Coyle, J., Black, B.E., Rekosh, D., Hammarskjold, M.L., and Paschal, B.M. (2001). RNA export mediated by tap involves NXT1-dependent interactions with the nuclear pore complex. The Journal of biological chemistry 276, 44953–44962.

Lim, A.K., and Kai, T. (2007). Unique germ-line organelle, nuage, functions to repress selfish genetic elements in Drosophila melanogaster. Proceedings of the National Academy of Sciences of the United States of America 104, 6714–6719.

Lund, E., Guttinger, S., Calado, A., Dahlberg, J.E., and Kutay, U. (2004). Nuclear export of microRNA precursors. Science 303, 95–98.

Lund, M.K., and Guthrie, C. (2005). The DEAD-box protein Dbp5p is required to dissociate Mex67p from exported mRNPs at the nuclear rim. Molecular cell 20, 645–651.

Luo, M.L., Zhou, Z., Magni, K., Christoforides, C., Rappsilber, J., Mann, M., and Reed, R. (2001). Pre-mRNA splicing and mRNA export linked by direct interactions between UAP56 and Aly. Nature 413, 644–647.

Mahat, D.B., Kwak, H., Booth, G.T., Jonkers, I.H., Danko, C.G., Patel, R.K., Waters, C.T., Munson, K., Core, L.J., and Lis, J.T. (2016). Base-pair-resolution genome-wide mapping of active RNA polymerases using precision nuclear run-on (PRO-seq). Nature protocols 11, 1455–1476.

Mahowald, A.P. (1971). Polar granules of Drosophila. 3. The continuity of polar granules during the life cycle of Drosophila. J Exp Zool 176, 329–343.

Malone, C.D., Brennecke, J., Dus, M., Stark, A., McCombie, W.R., Sachidanandam, R., and Hannon, G.J. (2009). Specialized piRNA pathways act in germline and somatic tissues of the Drosophila ovary. Cell 137, 522–535.

Meignin, C., and Davis, I. (2008). UAP56 RNA helicase is required for axis specification and cytoplasmic mRNA localization in Drosophila. Developmental biology 315, 89–98.

Mohn, F., Handler, D., and Brennecke, J. (2015). Noncoding RNA. piRNA-guided slicing specifies transcripts for Zucchini-dependent, phased piRNA biogenesis. Science 348, 812–817.

Mohn, F., Sienski, G., Handler, D., and Brennecke, J. (2014). The rhino-deadlock-cutoff complex licenses noncanonical transcription of dual-strand piRNA clusters in Drosophila. Cell 157, 1364–1379.

Morlan, J.D., Qu, K., and Sinicropi, D.V. (2012). Selective depletion of rRNA enables whole transcriptome profiling of archival fixed tissue. PloS one 7, e42882.

Ni, J.Q., Zhou, R., Czech, B., Liu, L.P., Holderbaum, L., Yang-Zhou, D., Shim, H.S., Tao, R., Handler, D., Karpowicz, P., et al. (2011). A genome-scale shRNA resource for transgenic RNAi in Drosophila. Nature methods 8, 405–407.

Nott, T.J., Petsalaki, E., Farber, P., Jervis, D., Fussner, E., Plochowietz, A., Craggs, T.D., Bazett-Jones, D.P., Pawson, T., Forman-Kay, J.D., et al. (2015). Phase transition of a disordered nuage protein generates environmentally responsive membraneless organelles. Molecular cell 57, 936–947.

Ozata, D.M., Gainetdinov, I., Zoch, A., O’Carroll, D., and Zamore, P.D. (2018). PIWI-interacting RNAs: small RNAs with big functions. Nature reviews Genetics.

Pan, J., Eckardt, S., Leu, N.A., Buffone, M.G., Zhou, J., Gerton, G.L., McLaughlin, K.J., and Wang, P.J. (2009). Inactivation of Nxf2 causes defects in male meiosis and age-dependent depletion of spermatogonia. Developmental biology 330, 167–174.

Pane, A., Jiang, P., Zhao, D.Y., Singh, M., and Schupbach, T. (2011). The Cutoff protein regulates piRNA cluster expression and piRNA production in the Drosophila germline. The EMBO journal 30, 4601–4615.

Patro, R., Duggal, G., Love, M.I., Irizarry, R.A., and Kingsford, C. (2017). Salmon provides fast and bias-aware quantification of transcript expression. Nature methods 14, 417–419.

Pimentel, H., Bray, N.L., Puente, S., Melsted, P., and Pachter, L. (2017). Differential analysis of RNA-seq incorporating quantification uncertainty. Nature methods 14, 687–690.

Port, F., Chen, H.M., Lee, T., and Bullock, S.L. (2014). Optimized CRISPR/Cas tools for efficient germline and somatic genome engineering in Drosophila. Proceedings of the National Academy of Sciences of the United States of America 111, E2967–2976.

Preibisch, S., Saalfeld, S., Schindelin, J., and Tomancak, P. (2010). Software for bead-based registration of selective plane illumination microscopy data. Nature methods 7, 418–419.

Raney, B.J., Dreszer, T.R., Barber, G.P., Clawson, H., Fujita, P.A., Wang, T., Nguyen, N., Paten, B., Zweig, A.S., Karolchik, D., et al. (2013). Track data hubs enable visualization of user-defined genome-wide annotations on the UCSC Genome Browser. Bioinformatics.

RCoreTeam (2018). R: A Language and Environment for Statistical Computing.

Reed, R., and Hurt, E. (2002). A conserved mRNA export machinery coupled to pre-mRNA splicing. Cell 108, 523–531.

Ren, Y., Schmiege, P., and Blobel, G. (2017). Structural and biochemical analyses of the DEAD-box ATPase Sub2 in association with THO or Yra1. Elife 6.

Schindelin, J., Arganda-Carreras, I., Frise, E., Kaynig, V., Longair, M., Pietzsch, T., Preibisch, S., Rueden, C., Saalfeld, S., Schmid, B., et al. (2012). Fiji: an open-source platform for biological-image analysis. Nature methods 9, 676–682.

Senti, K.A., Jurczak, D., Sachidanandam, R., and Brennecke, J. (2015). piRNA-guided slicing of transposon transcripts enforces their transcriptional silencing via specifying the nuclear piRNA repertoire. Genes & development 29, 1747–1762.

Shpiz, S., Ryazansky, S., Olovnikov, I., Abramov, Y., and Kalmykova, A. (2014). Euchromatic Transposon Insertions Trigger Production of Novel Pi- and Endo-siRNAs at the Target Sites in the Drosophila Germline. PLoS genetics 10, e1004138.

Siomi, M.C., Sato, K., Pezic, D., and Aravin, A.A. (2011). PIWI-interacting small RNAs: the vanguard of genome defence. Nature reviews Molecular cell biology 12, 246–258.

Smyth, G.K. (2004). Linear models and empirical bayes methods for assessing differential expression in microarray experiments. Statistical applications in genetics and molecular biology 3, Article3.

Strasser, K., and Hurt, E. (2001). Splicing factor Sub2p is required for nuclear mRNA export through its interaction with Yra1p. Nature 413, 648–652.

Tokuyasu, K.T. (1973). A technique for ultracryotomy of cell suspensions and tissues. The Journal of cell biology 57, 551–565.

Venken, K.J., Carlson, J.W., Schulze, K.L., Pan, H., He, Y., Spokony, R., Wan, K.H., Koriabine, M., de Jong, P.J., White, K.P., et al. (2009). Versatile P[acman] BAC libraries for transgenesis studies in Drosophila melanogaster. Nature methods 6, 431–434.

Vizcaino, J.A., Csordas, A., del-Toro, N., Dianes, J.A., Griss, J., Lavidas, I., Mayer, G., Perez-Riverol, Y., Reisinger, F., Ternent, T., et al. (2016). 2016 update of the PRIDE database and its related tools. Nucleic acids research 44, D447–456.

Wang, L., Dou, K., Moon, S., Tan, F.J., and Zhang, Z.Z. (2018). Hijacking Oogenesis Enables Massive Propagation of LINE and Retroviral Transposons. Cell 174, 1082–1094 e1012.

Webster, A., Li, S., Hur, J.K., Wachsmuth, M., Bois, J.S., Perkins, E.M., Patel, D.J., and Aravin, A.A. (2015). Aub and Ago3 Are Recruited to Nuage through Two Mechanisms to Form a Ping-Pong Complex Assembled by Krimper. Molecular cell 59, 564–575.

Wehr, K., Swan, A., and Schupbach, T. (2006). Deadlock, a novel protein of Drosophila, is required for germline maintenance, fusome morphogenesis and axial patterning in oogenesis and associates with centrosomes in the early embryo. Developmental biology 294, 406–417.

Wickham, H. (2016). ggplot2: Elegant Graphics for Data Analysis (Springer-Verlag New York).

Wickham, H. (2017). scales: Scale Functions for Visualization.

Yang, J., Bogerd, H.P., Wang, P.J., Page, D.C., and Cullen, B.R. (2001). Two closely related human nuclear export factors utilize entirely distinct export pathways. Molecular cell 8, 397–406.

Yi, R., Qin, Y., Macara, I.G., and Cullen, B.R. (2003). Exportin-5 mediates the nuclear export of pre-microRNAs and short hairpin RNAs. Genes & development 17, 3011–3016.

Zhang, F., Wang, J., Xu, J., Zhang, Z., Koppetsch, B.S., Schultz, N., Vreven, T., Meignin, C., Davis, I., Zamore, P.D., et al. (2012). UAP56 couples piRNA clusters to the perinuclear transposon silencing machinery. Cell 151, 871–884.

Zhang, G., Tu, S., Yu, T., Zhang, X.O., Parhad, S.S., Weng, Z., and Theurkauf, W.E. (2018). Co-dependent Assembly of Drosophila piRNA Precursor Complexes and piRNA Cluster Heterochromatin. Cell reports 24, 3413–3422 e3414.

Zhang, Z., Wang, J., Schultz, N., Zhang, F., Parhad, S.S., Tu, S., Vreven, T., Zamore, P.D., Weng, Z., and Theurkauf, W.E. (2014). The HP1 homolog rhino anchors a nuclear complex that suppresses piRNA precursor splicing. Cell 157, 1353–1363.

